# Firing-Rate Based Network Modeling of the dLGN Circuit: Effects of Cortical Feedback on Spatiotemporal Response Properties of Relay Cells

**DOI:** 10.1101/246140

**Authors:** Milad Hobbi Mobarhan, Geir Halnes, Pablo Martínez-Cañada, Torkel Hafting, Marianne Fyhn, Gaute T. Einevoll

## Abstract

Visual signals originating in the retina pass through the dorsal geniculate nucleus (dLGN), the visual part of thalamus, on the way to the visual cortex. This is however not a simple feedforward flow of information: there is a significant feedback from cortical cells back to both relay cells and interneurons in the dLGN. Despite four decades of experimental and theoretical studies, the functional role of this feedback is still debated. Here we use a firing-rate model, the extended difference-of-gaussians (eDOG) model, to explore cortical feedback effects on visual responses of dLGN relay cells. For this model the responses are found by direct evaluation of two- or three-dimensional integrals allowing for fast and comprehensive studies of putative effects of different candidate organizations of the cortical feedback. Our analysis identifies a special mixed configuration of excitatory and inhibitory cortical feedback which seems to best account for available experimental data. This configuration consists of a slow (long-delay) and spatially widespread inhibitory feedback, combined with a fast (short-delayed) and spatially narrow excitatory feedback, where (iii) the excitatory/inhibitory ON-ON connections are accompanied respectively by inhibitory/excitatory OFF-ON connections, i.e. following a phase-reversed arrangement. The recent development of optogenetic and pharmacogenetic methods has provided new tools for more precise manipulation and investigation of the thalamocortical circuit, in particular for mice. Such data will expectedly allow the eDOG model to be better constrained by data from specific animal model systems than has been possible until now for cat. We have therefore made the Python tool `pyLGN` which allows for easy adaptation of the eDOG model to new situations.

**Author Summary:** On route from the retina to primary visual cortex, visually evoked signals have to pass through the dorsal lateral geniculate nucleus (dLGN). However, this is not an exclusive feed forward flow of information as feedback exists from neurons in the cortex back to both relay cells and interneurons in the dLGN. The functional role of this feedback remains mostly unresolved. Here, we use a firing-rate model, the extended difference-of-gaussians (eDOG) model, to explore cortical feedback effects on visual responses of dLGN relay cells. Our analysis indicates that a particular mix of excitatory and inhibitory cortical feedback agrees best with available experimental observations. In this configuration ON-center relay cells receive both excitatory and (indirect) inhibitory feedback from ON-center cortical cells (ON-ON feedback) where the excitatory feedback is fast and spatially narrow while the inhibitory feedback is slow and spatially widespread. In addition to the ON-ON feedback, the connections are accompanied by OFF-ON connections following a so-called phase-reversed (push-pull) arrangement. To facilitate further applications of the model, we have made the Python tool `pyLGN` which allows for easy modification and evaluation of the a priori quite general eDOG model to new situations.

## 1 Introduction

Visual signals originating in the retina pass through the dorsal geniculate nucleus (dLGN), the visual part of thalamus, on the way to the visual cortex. This is however not a simple feedforward flow of information, as there is a significant feedback from primary visual cortex back to dLGN. Cortical cells feed back to both relay cells and interneurons in the dLGN, and also to cells in the thalamic reticular nucleus (TRN) which in turn provide feedback to dLGN cells [1,2]. In the last four decades numerous experimental studies have provided insight into the potential roles of this feedback in modulating the transfer of visual information in the dLGN circuit [3–19]. Cortical feedback has been observed to switch relay cells between tonic and burst response modes [20, 21], increase the center-surround antagonism of relay cells [16, 17, 22, 23], and synchronize the firing patterns of groups of such cells [10, 13]. However, the functional role of cortical feedback is still debated [1, 24–30].

Several studies have used computational modeling to investigate cortical feedback effects on spatial and/or temporal visual response properties of dLGN cells [31–39]. These have typically involved numerically extensive dLGN network simulations based on spiking neurons [31–33, 35, 39] or models where each neuron is represented as individual firing-rate unit [36,37]. This is not only computationally cumbersome, but the typically large number of model parameters in these comprehensive network models also makes a systematic exploration of the model behaviour very difficult.

In the present study we instead use a firing-rate based model, the extended *difference-of-gaussians* (eDOG) model [40], to explore putative cortical feedback effects on visual responses of dLGN relay cells. A main advantage with this model is that visual responses are found from direct evaluation of two-dimensional or three-dimensional integrals in the case of static or dynamic (i.e., movie) stimuli, respectively. This computational simplicity allows for fast and comprehensive study of putative effects of different candidate organizations of the cortical feedback. Taking advantage of the computational efficiency of the eDOG model, we here explore effects of direct excitatory and indirect inhibitory feedback effects (via dLGN interneurons and TRN neurons) on spatiotemporal responses of dLGN relay cells. In particular we investigate effects of (i) different spatial spreads of corticothalamic feedback and (ii) different corticothalamic propagation delays.

Our analysis suggests that a particular mix of excitatory and inhibitory cortical feedback agrees best with available experimental observations. In this configuration an ON-center relay cell receives feedback from ON-center cortical cells (ON-ON feedback), consisting of a slow (long-delay) and spatially widespread inhibitory feedback combined with a fast (short-delay) and spatially narrow excitatory feedback. Here the inhibitory and excitatory ON-ON feedback connections are accompanied by excitatory and inhibitory OFF-ON connections, respectively, following a phase-reversed arrangement [39]. For one this feedback organization accounts for the feedback-induced enhancement of center-surround antagonism of relay cells as observed in experiments [16,17,22,23,39]. Further, it seems well suited to dynamically modulate both the center-surround suppression and spatial resolution, for example, to adapt to changing light conditions [41].

Morever, a longer thalamocortical loop time of ON-ON inhibitory feedback loop compared to ON-ON excitatory feedback may contribute to temporal decorrelation of natural stimuli [42], an operation that has been observed accomplished at the level of dLGN in the early visual pathway [43]. At the same time, the rapid excitatory feedback may contribute to linking stimulus features by synchronising firing of neighbouring relay cells [10, 19].

Previous experimental studies have focused on cat, monkey and ferret dLGN, and the present model was adapted to neurobiological findings from cat. However, the last years have seen a surge of interest in mouse visual system, where new optogenetic and pharmacogenetic methods provide new tools for precise manipulation of identified neurons in the thalamocortical circuit [44–49]. Such data will expectedly allow for a detailed adaptation of the eDOG model to rodent dLGN, likely much better constrained by biological findings than what has been possible until now for cat. To facilitate this we have made the Python tool `pyLGN` (http://pylgn.rtfd.io) which allows for easy modification and evaluation of the eDOG model to new situations.

## 2 Materials and Methods

### 2.1 Spatiotemporal receptive fields

Spike responses of neurons in the early visual pathway are most commonly described in terms of *receptive fields*, the most common measure of spiking activity in the visual system. Mathematically, the spatiotemporal receptive field is defined by an impulse-response function *W* (**r**, *t*). This function describes the firing-rate response to a tiny (*δ*-function) spot positioned at **r** = 0 which is on for a very short time (*δ*-function) at *t* = 0. If linearity is assumed, the response to any stimulus *S*(**r**, *t*) can be found by convolving the impulse-response function with the stimulus [40,50–53]:

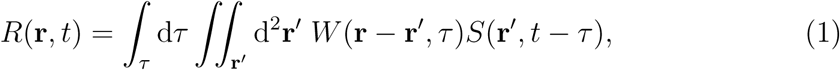

or written more compactly

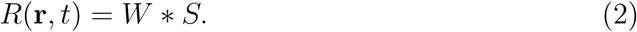

Here *S*(**r**, *t*) is a spatiotemporal stimulus function describing, e.g., the light intensity on a screen as a function of time and position. *R*(**r**, *t*) is the response of a neuron with its receptive-field center at **r**. The spatial integral goes over the whole visual field, i.e., over all two-dimensional space. For mathematical convenience we have chosen the temporal integral to go from *τ* = −∞ to +∞. Since a stimulus input cannot affect the response in the past, it then follows that *W*(**r**, *τ* < 0) = 0.

In Fourier space the convolution in Eq. (2) corresponds to a product

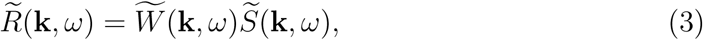

where
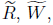
 and 
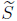
 are the Fourier transforms of the neural response *R*, the impulse-response function *W*, and the stimulus *S*, respectively. The tilde symbol 
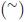
 will be used to denote the Fourier transform of any function throughout this paper. The function argument **k** is the wave vector which is related to the spatial frequency 
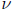
 via 
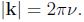
 Correspondingly, the angular frequency *ω* is related to the temporal frequency *f* via *ω* = 2*π f*. With 
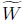
 and 
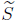
 known, the neural response can thus always be found by an inverse Fourier transform 
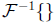
, which entails an integral over temporal and spatial frequencies

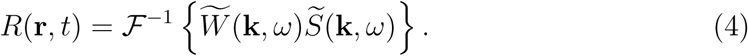

The response model in Eq. (1) is an example of a *descriptive* model where the purpose is to summarize experimental data compactly in a mathematical form [52–54]. Here the aim is to find an appropriate impulse-response function, i.e., spatiotemporal receptive-field function, that describes the measured neural response to different visual stimuli [51]. With this approach, however, limited insight is gained into how the neurons and neural circuitry in the early visual system provide such a receptive field. To address this question a *mechanistic* receptive-field model is needed. (For a discussion of the difference between descriptive and mechanistic models in visual neuroscience, see, for example [54,55].)

### 2.2 Mechanistic receptive-field models

In mechanistic LGN-circuit models the input from retinal ganglion cells have been described by descriptive models, see, e.g., [36,37,40,53,56]. Likewise, in the present eDOG model the input from retinal ganglion cells is represented by the descriptive impulse-response function (Eq. (1)). Here a square grid of retinal ganglion cells with identical, spatially-localised receptive fields are considered (see Fig. 1). The activity, i.e., firing rates, of the neurons on the retinal ganglion cell layer then serves as input to the dLGN relay cell layer. This is represented by a spatiotemporal coupling-kernel function *K*_RG_, which reflects the direct synaptic input from retinal ganglion cells to dLGN relay cells. The coupling kernel, which is analogous to the descriptive impulse-response function in Eq. (1), is assumed to only depend on the relative distance between the cells in the visual field [53].

**Fig 1.**
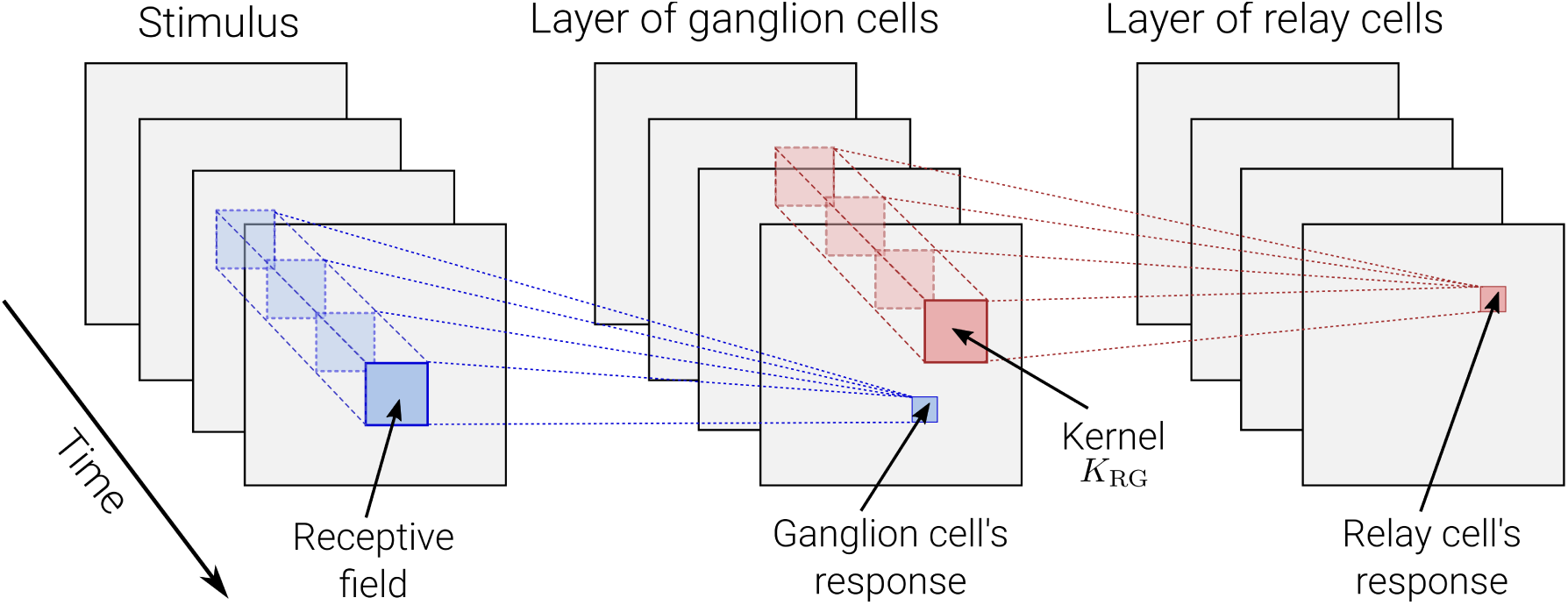
Illustration of mechanistic model. A dense and evenly distributed layer of retinal ganglion cells with identical response properties are activated by the visual stimulus according to their receptive fields. This creates a pattern of neural activity for the layer of ganglion cells, acting as input for a similar layer of dLGN relay cells. Relay cells are connected to the ganglion cells via a spatiotemporal coupling-kernel function *K*_RG_ which is assumed to only depend on the relative distance between the retinal ganglion cell and relay cell.

The response of a relay cell located at r is then given by [40,53]:

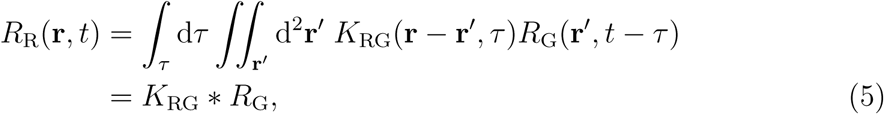

where *R*_R_ and *R*_G_ are the firing-rate responses of relay cells and ganglion cells, respectively. The coupling kernel 
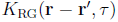
 denotes the strength with which the response of a ganglion cell, displaced by **r** − **r′** from the relay cell, at time *t* − *τ* influences the response of the latter at time *t*. Note that, 
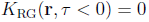
 due to causality.

In Fourier space the relationship in Eq. (5) can be written as

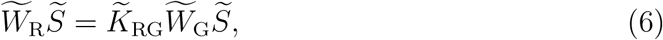

where we have used the general relationship in Eq. (4). The key point here is that a *descriptive* model for the relay-cell impulse-response function 
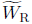
 now has a *mechanistic* interpretation. This relation is given as the product of the impulse-response function 
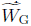
 of the retinal ganglion cells and the coupling kernel 
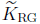
 from the former cell type to the latter.

In the eDOG model this approach is extended to include the various feedforward and feedback connections affecting the relay-cell response. The result is an expression for the relay-cell impulse-response function 
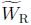
 in terms of the impulse-response function
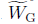
 of the retinal input and the coupling kernels connecting the neurons of the circuit. With such a mechanistic expression for 
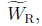
 the response to any visual stimulus can be computed by means of the inverse Fourier transform in Eq. (4).

### 2.3 Extended Difference-of-Gaussians (eDOG) model

Here we derive the impulse-response function for dLGN relay cells for the mechanistic eDOG model [40]. The complete circuit is shown in Fig. 2. In this figure each cell type correspond to a two-dimensional layer (or population) of identical cells.

**Fig 2.**
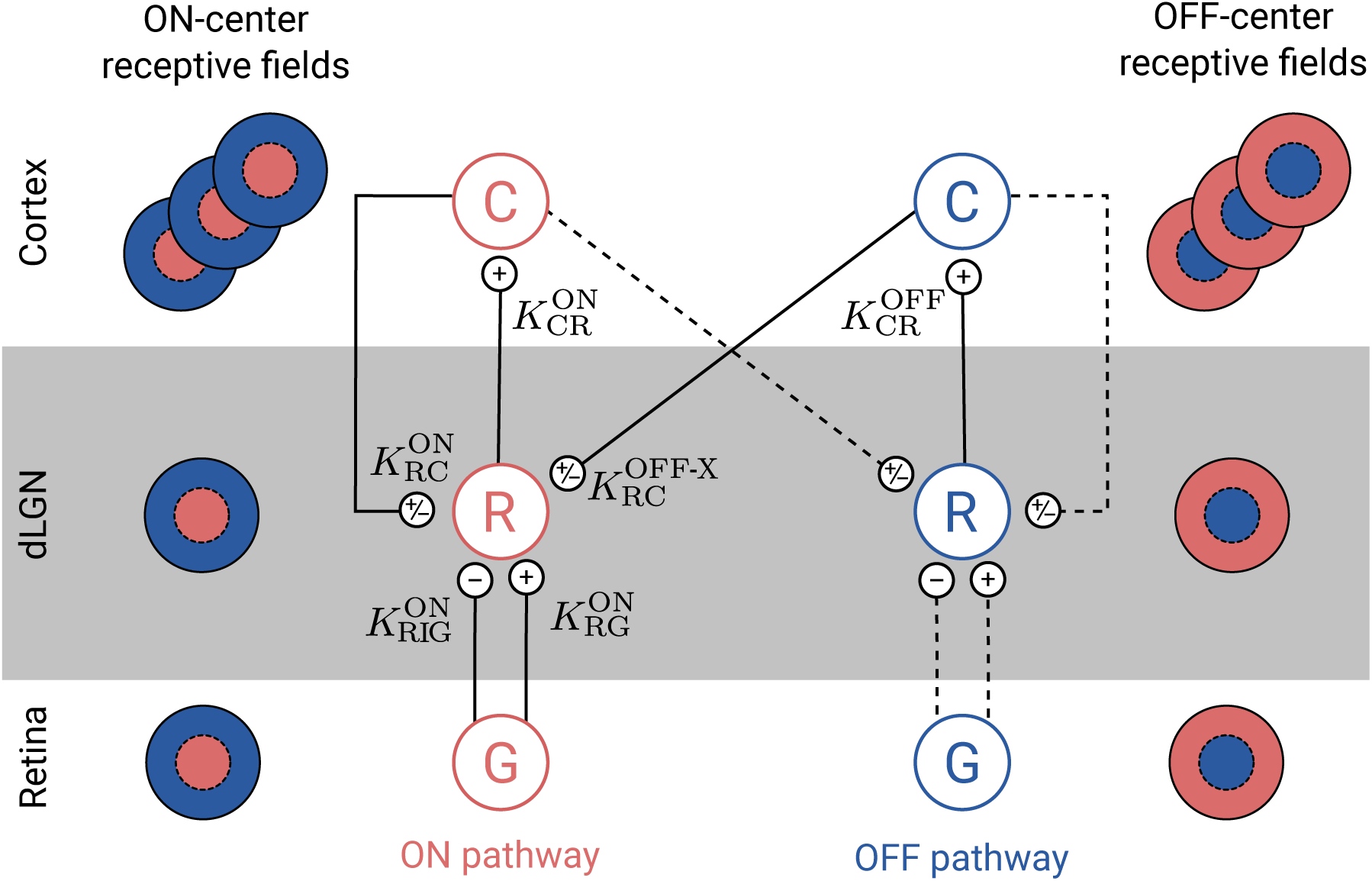
Schematic overview of the present eDOG model. Cell types are: retinal ganglion cells (G), dLGN relay cells (R), and cortical cells (C). Each cell type corresponds to a two-dimensional layer (or population) of identical cells (see Fig. 1). Note that only one cortical population is shown for each pathway even though an arbitrary number of cortical populations is considered. Unlike the feedforward projection, the feedback is cross-symmetry, i.e., the activity of ON-center relay cells are affected both by ON and OFF-center cortical cells. The OFF-center dLGN relay cells are assumed to receive the same input as the corresponding ON-center dLGN relay cells with opposite sign. Solid lines represent explicitly included connections in the eDOG model, while dashed lines represent connections included implicitly.

We will in the following focus on the dLGN relay cells with ON symmetry, but a similar model can be constructed for OFF-symmetry cells. These neurons receive feedforward excitation and indirect feedforward inhibition (via intrageniculate interneurons) from ON-center ganglion cells in retina. The relay cells further receive cortical feedback from both cortical ON cells and cortical OFF cells.

#### 2.3.1 Feedforward input from retina

With indirect feedforward inhibition included in addition to the direct feedforward excitation, the expression in Eq. (5) generalizes to

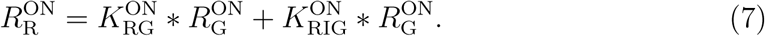

Here 
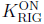
 is a spatiotemporal coupling-kernel representing the indirect feedforward inhibition from retinal ganglion cells onto relay cells via intrageniculate interneurons.

In Fourier space this gives a simple expression for the relay-cell impulse-response function, i.e.,

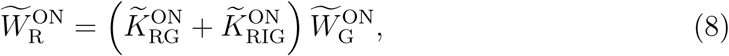

where we have used that 
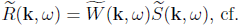
. Eq. (3).

#### 2.3.2 Feedback from cortex

Next we add effects from cortical feedback onto the relay cell. This cortical feedback can be both excitatory and inhibitory. The excitatory feedback corresponds to direct projections from cortical cells onto relay cells. The inhibitory feedback corresponds to indirect inhibitory action on relay cells mediated by cortical projections onto inhibitory TRN and intrageniculate interneurons. Further, unlike the feedforward projection, the feedback is cross-symmetry, i.e., the activity of ON relay cells are affected both by ON and OFF cortical cells.

In the eDOG model cortical ON and OFF cells are assumed to be driven solely by ON and OFF relay cells, respectively. As the corticogeniculate feedback comes from orientation-tuned cells in layer 6 in cortex, we include a set of N mutually uncoupled, orientation-selective cortical populations 
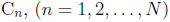
 for both the ON and OFF pathways. Each population C_*n*_ responds preferably to stimuli (bars, gratings) with orientation *θ_n_*. In Fig. 2 only a single cortical population is shown for each pathway even though an arbitrary number of *N* cortical populations can be considered.

Cortical cells are known to exhibit substantial non-linearities when responding to visual stimuli, and here the response is modeled via a static non-linear function acting on a linearly filtered input [52,57]. More specifically we express the response of the ON or OFF cortical population C_*n*_ by

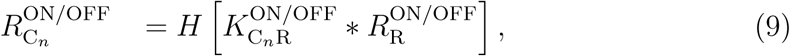

where 
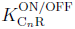
is the feedforward kernel between the relay cells and the cortical cells in population C_*n*_. Further, the half-wave rectification function *H*[*x*] = *xθ*(*x*) is used to enforce non-negative firing rates [58], where *θ*(*x*) is the Heaviside step function.

We further assume the input to cortical OFF cells to be the negative of the one for the ON cells [54]. That is

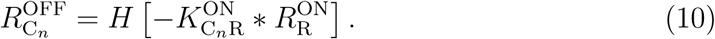

Finally, the feedback cross-connection (OFF to ON) is assumed to be phase-reversed compared to the same-sign feedback (ON to ON) [40]:

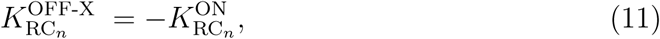

where 
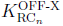
 is the cross-coupling feedback from cortical OFF cells onto relay ON cells. In other words, we assume the effect of ON-center and OFF-center cortical cells to be the opposite of each other. However, we do not make any specific assumptions on whether, say, the excitatory or inhibitory feedback is driven by ON-center or OFF-center cortical cells [40].

With the three assumptions in Eqs. (9)–(11), the total input to the ON dLGN relay cell is found to be [40,54]

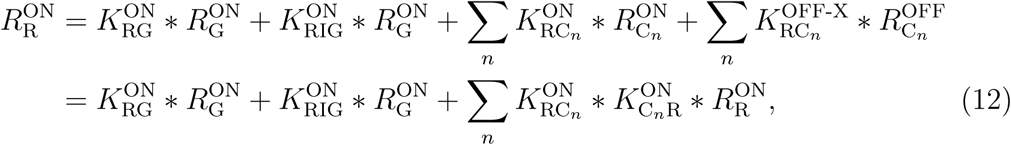

where we have used the mathematical identity: 
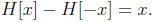

In Fourier space we thus have

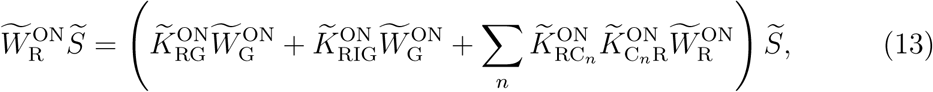

and in analogy with Eq. (8) we find after some simple algebra

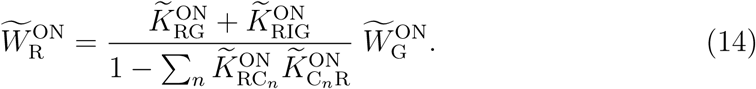

In this expression the direct feedforward excitation and the indirect feedforward inhibition via interneurons are represented by the first and second terms in the numerator, respectively. The feedback effects are accounted for in the denominator.

The general mathematical expression in Eq. (14) for the (Fourier transformed) impulse-response function for the relay cells is the main feature for the eDOG model [40]. The model provides an analytical formula for (linear) impulse-response function for relay cells, despite the non-linearity of the response of the cortical cells providing the feedback. The simulator presented in this paper uses this expression as basis to compute the impulse-response function and the spatiotemporal responses for user-defined kernels and input stimuli. Once the explicit form of the kernels in Eq. (14) are defined, the response of the relay cells to arbitrary stimuli can be calculated using Eq. (4).

In the next subsections we describe the choices made in this paper for (i) the descriptive spatiotemporal receptive-field function for the retinal input 
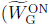
 in Eq. (14)), (ii) the various mechanistic coupling kernels inside the dLGN circuit ( 
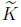
 in Eq. (14)), and (iii) the visual stimulus 
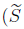
 in Eq. (4)). The coupling kernels are

assumed to be space-time separable (e.g., 
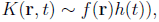
 but space-time coupled kernels can equally be used in the eDOG-model. The same applies to the choice of the receptive-field function of the retinal input [59,60]. For presentational simplicity, we will focus on the ON-pathway and skip the ON-superscript, but an analogous model is equally applicable for the OFF pathway.

#### 2.3.3 Impulse-response function of input from retinal ganglion cells

The impulse-response function of the retinal input is modeled as a product of a spatial part *F*(**r**) and temporal part *H*(*t*). The spatial part is described by means of the difference-of-Gaussians (DOG) model [61]:

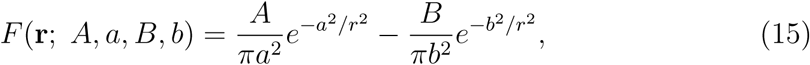

where the first and second term correspond to the center and surround contribution, respectively. Further, *A* and *B* (defined to be positive) are the strengths of the center and surround, and a and b are the corresponding width parameters. In the present paper we have used parameters extracted from fitting the function to retinal-input responses to flashing circular spots [56].

The temporal part of the impulse response of the retinal input is modeled as a biphasic temporal function [37,54]:

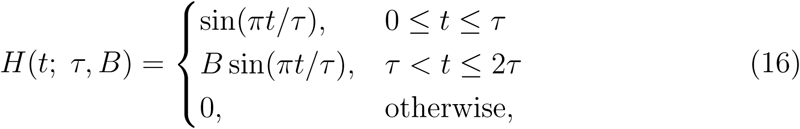

where *B* is the weight for the second phase, and *τ* is the duration of each phase. The same parameter values as in [54] has been used, which correspond to the mean of the range of values reported by [62].

For an illustration of the shapes of the spatial and temporal impulse-response function, see Fig. 3.

**Fig 3.**
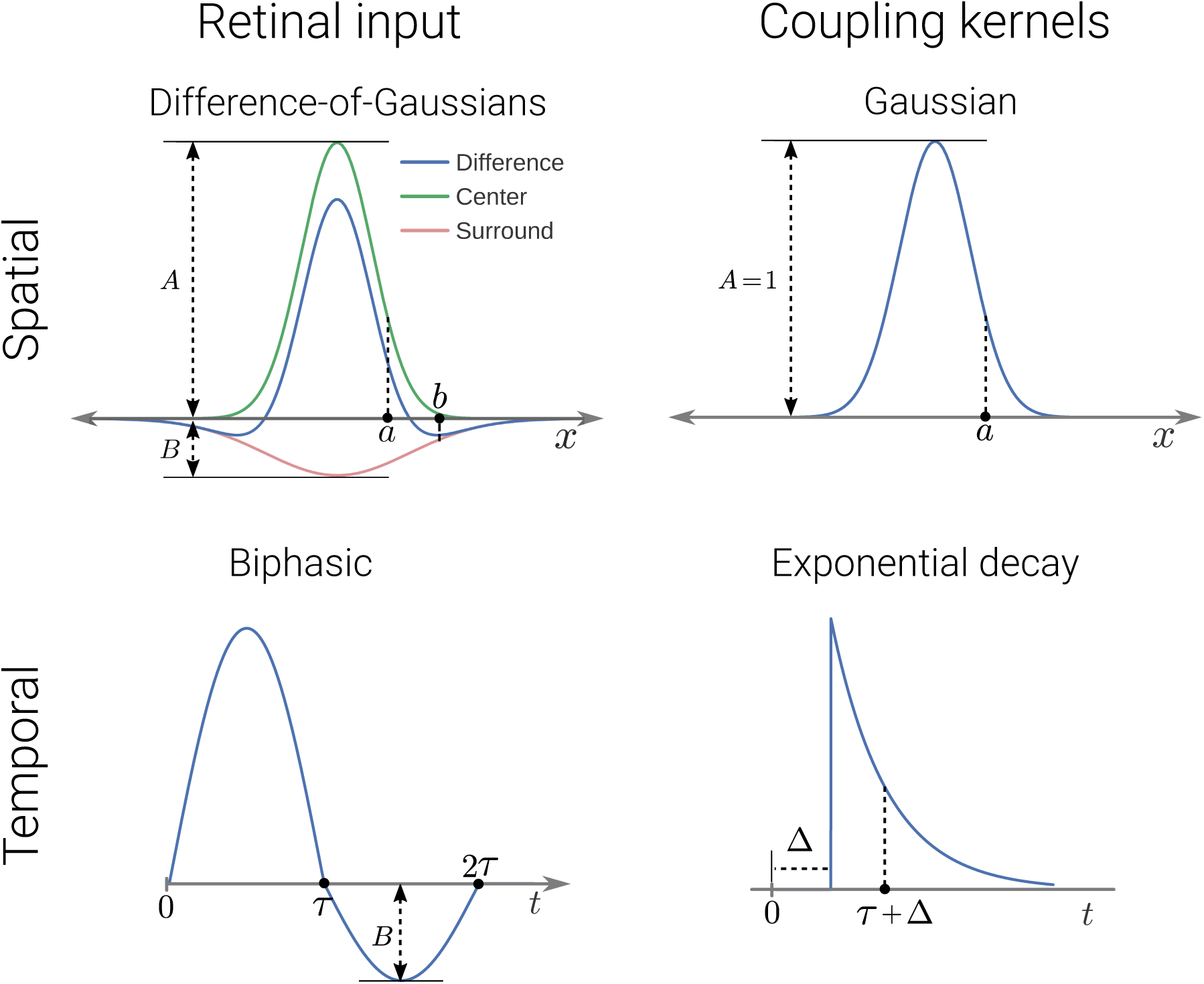
The shape of the spatial and temporal part of the receptive-field function for the retinal input and the connectivity kernels. Left panel shows the receptive-field function for the retinal input to dLGN cells and right panel shows the connectivity kernels. The spatial functions are shown as one-dimensional plot, although they are (circularly symmetric) two-dimensional functions.

#### 2.3.4 Coupling kernels inside dLGN circuit

The kernels *K*(**r**, *t*) are considered to have separable space-time parts, i.e.,

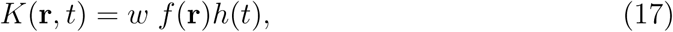

where *f* and *h* are normalised spatial and temporal parts, respectively, and *w* is the connection weight of the kernel. The latter is positive for excitatory synaptic connections and negative for inhibitory connections. The normalization implies that 
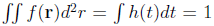
 where the integrals go over all visual (two-dimensional) space and all times, respectively.

The spatiotemporal coupling-kernels in the circuit, reflecting how the firing in one type of cell affects the firing in another type of cell through their direct synaptic connections, have not been systematically mapped out. However, a key design principle of the early visual pathway is retinotopy, i.e., that neurons representing neighboring positions in the visual field also are neighbors inside the retina, dLGN, and visual cortex. This implies that the coupling kernels are spatially confined. In this paper we describe the shape of spatial kernels using the mathematically convenient Gaussian function:

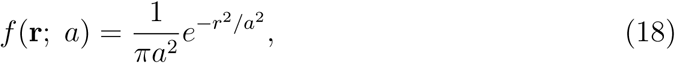

where *a* is the width parameter.

The temporal part of the kernels is modeled as (delayed) exponential decay in accordance to previous modeling studies [53,54]:

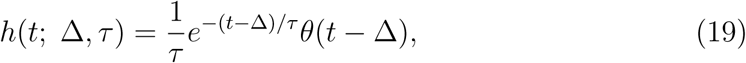

where *τ* is the time constant, and Δ corresponds to a combined axonal and synaptic time delay. For an illustration of the shapes of spatial and temporal part of the coupling kernels, see Fig. 3.

We next describe the kernel parameters used for the circuit coupling. A detailed list of these parameters is given in Table 1.

**Table 1.** List of kernel parameters. *W*_G_ is the impulse-response function of ganglion cells, *K*_RG_ and *K*_RIG_ are the excitatory and inhibitory feedforward kernels, respectively. 
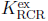
 and 
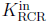
 are ON-ON excitatory and inhibitory thalamo-cortico-thalamic kernels, respectively. 
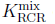
 enotes the mixed ON-ON feedback kernel, consisting of an excitatory and an inhibitory term. *F* represents the DOG function, *f* represents the Gaussian function, *H* represents the biphasic temporal function, and *h* represents the delayed decaying exponential function. The width parameters in spatial functions are given in units of degree, while the temporal parameters are in units of ms. In the present example applications we have kept the time constant *τ* fixed at 5 ms (comparable to what, e.g., was found in [75]), while the temporal delay parameters Δ have been varied in a range of 5-30 ms. ^†^ denotes the default values for parameters that have been varied.

| Kernel | Weight | Spatial | Temporal |
| --- | --- | --- | --- |
| $W_G$ | | $F(\mathbf{r}; A_G=1, a_G=0.62, B_G=0.85, b_G=1.26)$ | $H(t; \tau_G = 42.5, B = 0.38)$ |
| $K_{RG}$ | $w_{RG}=1$ | $f(\mathbf{r}; a_{RG} = 0.1)$ | $h(t; \Delta_{RG} = 0, \tau_{RG} = 5)$ |
| $K_{RIG}$ | $w_{RIG}=-0.5^\dagger$ | $f(\mathbf{r}; a_{RIG} = 0.3^\dagger)$ | $h(t; \Delta_{RIG} = 3^\dagger, \tau_{RIG} = 5)$ |
| $K_{RCR}^{\text{ex}}$ | $w_{RCR}^{\text{ex}}=0.5^\dagger$ | $f(\mathbf{r}; a_{RCR}^{\text{ex}} = 0.83^\dagger)$ | $h(t; \Delta_{RCR}^{\text{ex}} = [5, 30], \tau_{RCR}^{\text{ex}} = 5)$ |
| $K_{RCR}^{\text{in}}$ | $w_{RCR}^{\text{in}}=-0.5^\dagger$ | $f(\mathbf{r}; a_{RCR}^{\text{in}} = 0.83^\dagger)$ | $h(t; \Delta_{RCR}^{\text{in}} = [5, 30], \tau_{RCR}^{\text{in}} = 5)$ |
| $K_{RCR}^{\text{mix}}$ | $w_{RCR}^{\text{ex}}=0.3^\dagger$ | $f(\mathbf{r}; a_{RCR}^{\text{ex}} = 0.1^\dagger)$ | $h(t; \Delta_{RCR}^{\text{ex}} = [5, 30], \tau_{RCR}^{\text{ex}} = 5)$ |
| | $w_{RCR}^{\text{in}}=-0.6^\dagger$ | $f(\mathbf{r}; a_{RCR}^{\text{in}} = 0.9^\dagger)$ | $h(t; \Delta_{RCR}^{\text{in}} = [5, 30], \tau_{RCR}^{\text{in}} = 5)$ |

Feedforward couplings. Relay cells in the cat appear to receive input from a single or a few retinal ganglion cells [63–69]. Further, the relay cells receive indirect feedforward inhibition via intrageniculate interneurons which in turn receive input from a few retinal ganglion cells [68,70]. Based on these observations and the known ‘retionotopographical’ organization of of the early visual pathway, we here use narrow Gaussian functions as coupling kernels between the retinal ganglion cells and dLGN relay cells [40,53]. We assume a larger width parameter for the feedforward inhibitory coupling kernel compared to the excitatory kernel [56], reflecting the observed larger receptive field in intrageniculate interneurons compared to both retinal ganglion cells and relay cells [68].

Feedback coupling. The net feedback coupling from cortex to relay cells are determinined by two factors: (i) the spatiotemporal response of the cortical cells providing the feedback, and (ii) the spatiotemporal feedback coupling kernels from cortical to LGN cells. The receptive fields of simple cortical cells arises primarily from convergent input from ON and OFF relay cells [71–73]. In order to model orientation-selective cortical populations, the thalamocortical kernels 
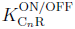
 in Eq. (9) must have an elongated shape. In [40] these kernels were, for example, modeled as elliptical Gaussians.

As seen in the denominator of Eq. (14), the total effect of cortical feedback is a sum over feedback contributions from all *n* populations, covering all orientation angles. Thus, the net feedback effect is expected to be essentially circularly symmetric [40]. The net effect of the cortical feedback from all ca n the model via a single circularly-symmetric coupling kernel 
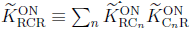
. As for the feedforward couplings, we for simplicity model the feedback coupling kernels as product of a Gaussian function of space (Eq. (18)) with a delayed exponentially-decaying temporal function (Eq. (19)).

The structure of the eDOG model is indifferent to whether the cortical feedback is excitatory, inhibitory, or even a mix of excitatory and inhibitory feedback. For excitatory feedback the weight parameter *w* in Eq. (17) is positive, while for inhibitory feedback it is negative. For mixed feedback the coupling kernel 
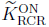
 consists of a sum of excitatory and inhibitory feedback terms. Note that in all cases the ON to ON couplings are accompanied by OFF to ON couplings with the opposite sign, i.e., a phase-reversed arrangement as described in Eqs. (10) and (11).

A few experiments give some hints about how the feedback may be organized. In [3] a center-surround feedback configuration was reported in cats where feedback was excitatory when the cortical and relay cell receptive field centers were close to each other and inhibitory when they were further apart. This observation was later supported by [16], where they found in primates a center-surround configuration for feedback, with a facilitatory bias to center and inhibitory surround (but see also [18]). Further, in [74] a particular cross-symmetry organization was observed where a same-symmetry inhibitory feedback was accompanied by an excitatory feedback with opposite symmetry, e.g., ON-ON inhibitory feedback accompanied by OFF-ON excitatory feedback.

In this paper we will study three different spatial organization of the cortical feedback as shown in the list below and illustrated in Fig. 4. In this list ON-ON refers to feedback from ON-center cortical cells to ON-center relay cells, while OFF-ON refers to feedback from OFF-center cortical cells to ON-center relay cells.

- ON-ON excitatory feedback (*K*^ex^_RCR_) combined with OFF-ON inhibitory feedback.
- ON-ON inhibitory feedback (*K*^in^_RCR_) combined with OFF-ON excitatory feedback.
- Mixed ON-ON excitatory and inhibitory feedback (*K*^mix^_RCR_). The OFF-ON feedback is also both excitatory and inhibitory.

**Fig 4.**
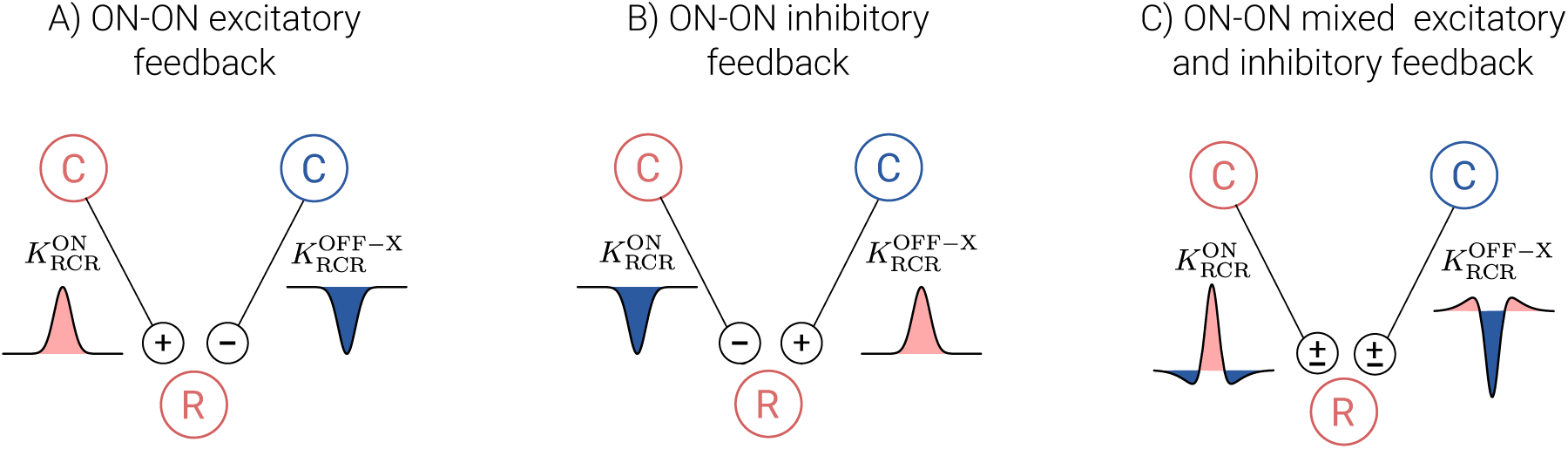
Spatial feedback configurations investigated in present study. The ON and OFF cells are marked with red and blue color, respectively. The spatial connectivity kernels are shown as one-dimensional plots where the fill color corresponds to the sign of the input (excitatory: red, inhibitory: blue).

Here, and in the following, the ON-superscript is skipped to simplify notation. The superscripts ‘ex’ (excitatory) and ‘in’ (inhibitory) refer to the sign of the ON-ON feedback, and the subscript ‘RCR’ refers to the complete thalamo-cortico-thalamic loop (relay → cortex → relay). These three scenarios are illustrated in Fig. 4. The second scenario corresponds to the configuration observed experimentally in [74], while the last configuration is inspired of the center-surround configuration suggested by data from [3,16]. For simplicity we will in the following refer to ON-ON excitatory feedback as just excitatory feedback and ON-ON inhibitory feedback as inhibitory feedback. It is then implicitly assumed that the influence from the OFF-ON feedback has the opposite sign.

The influence of each of these feedback con on the relay cell responses is investigated for a range of feedback strenghts *w*, width values *a* for the Gaussian functions (Eq. (18)), as well to temporal delays Δ of the delayed exponential functions (Eq. (19)).

#### 2.3.5 Visual stimuli

With the general eDOG relay-cell impulse-response function expression from Eq. (14), specified by the coupling kernels above, all that is needed to compute the relay-cell response by means of Eq. (4) is a mathematical expression for the stimulus *S*(**r**, *t*). The two main visual stimuli considered in the present work are (i) circular patch gratings and (ii) full-field gratings. For a full-field drifting grating, specified by **k**_g_ and *ω*_g_, the relay-cell response is essentially given by Fourier-transformed impulse response in Eq. (14) [40].

For a circular patch of drifting grating, the stimulus can be described mathematically as [40,76]

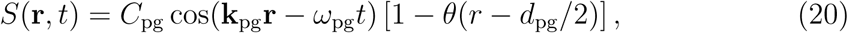

where **k**_pg_ and *ω*_pg_ are the wave vector and the angular frequency of the patch-grating, respectively, *d*_pg_ is the diameter of the patch-grating spot, and *C*_pg_ is a measure for the contrast of the grating. In all calculations presented in this paper *C*_pg_ = 1. Note that a static circular patch (spot) is obtained for **k**_pg_ = *ω*_pg_ = 0. In the limit *d*_pg_ → ∞ the Heaviside function in Eq. (20) is always zero, and we obtain the simple harmonic function representing a full-field grating.

In addition, natural stimulus (image and movie) is also used. The stimulus is then given as an array of numbers, and the Fourier transform of the stimulus is calculated numerically.

### 2.4 Implementation in pyLGN

In order to allow for easy exploration of the eDOG model and in particular effects of cortical feedback on relay-cell responses, we have developed an efficient, firing-rate based simulator of spatiotemporal responses in the early visual system. The simulator is named `pyLGN` and is written in Python. The design goals for `pyLGN` are to provide a software framework for studying the cortical feedback effects that is easy to use, extensible, and open. To facilitate usability, `pyLGN` has its own documentation page including installation instructions, several usage examples, and technical aspects (http://pylgn.rtfd.io). To achieve extensibility, object-oriented programming is used, making it possible for the user to define new connectivity kernels and input stimuli. Lastly, to support openness `pyLGN` is both open-source and multi-platform.

All calculations presented in this paper have been tracked using the Python software Sumatra [77], which is an automated tracking tool for computational simulations and analysis. The source code for all presented simulations is available at (https://github.com/miladh/edog-simulations).

## 3 Results

The result section is divided into two distinct parts. In the first part, results for the effects of cortical feedback on the spatial response properties of relay cells are presented (Sec. 3.1). The cortical feedback effects on temporal aspects are presented in the second part (Sec. 3.2).

### 3.1 Effect of cortical feedback on spatial properties

#### 3.1.1 Spatial receptive fields

We start our study of the dLGN network model by characterizing the effects of cortical feedback on the spatial aspects of the relay cell’s receptive field structure. From Eq. (14) we see that even with separable kernels the impulse-response function, in general, remains non-separable in space and time. However, with a static stimulus, the spatial response properties can be studied in isolation.

Mathematically, the Fourier transform 
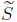
 for a static stimulus is 
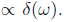
 The convolution in the response integral in Eq. (4) is then given by

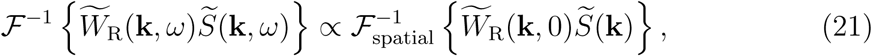

where 
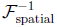
 is the spatial inverse Fourier transform, and 
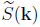
 is the Fourier transform of the spatial part of the stimulus.

Using the kernels shown in Table 1 we then find that the static relay-cell impulse response function 
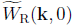
 is given by

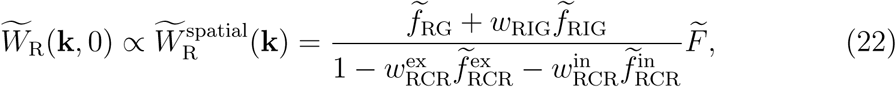

where we have used that 
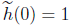
 and that 
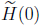
 is a constant. For simplicity will we hereafter refer t 
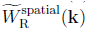
 as the spatial impulse-response function of relay cells. Examples of spatial receptive fields, found by an inverse Fourier transform of this function, is shown in Fig. 5. As seen in this figure the center-surround receptive field structure of the retinal ganglion cells is qualitatively preserved, in accordance with the notion that cortical feedback has a mainly modulatory effect on response properties of relay cells [78]. A close inspection of the right panel in the figure reveals that while the response at the receptive-field center (peak value) is increased for excitatory feedback, it is reduced for inhibitory feedback. The cortical feedback effects outside the receptive-field center, on the other hand, are less clear-cut for the examples in the figure.

**Fig 5.**
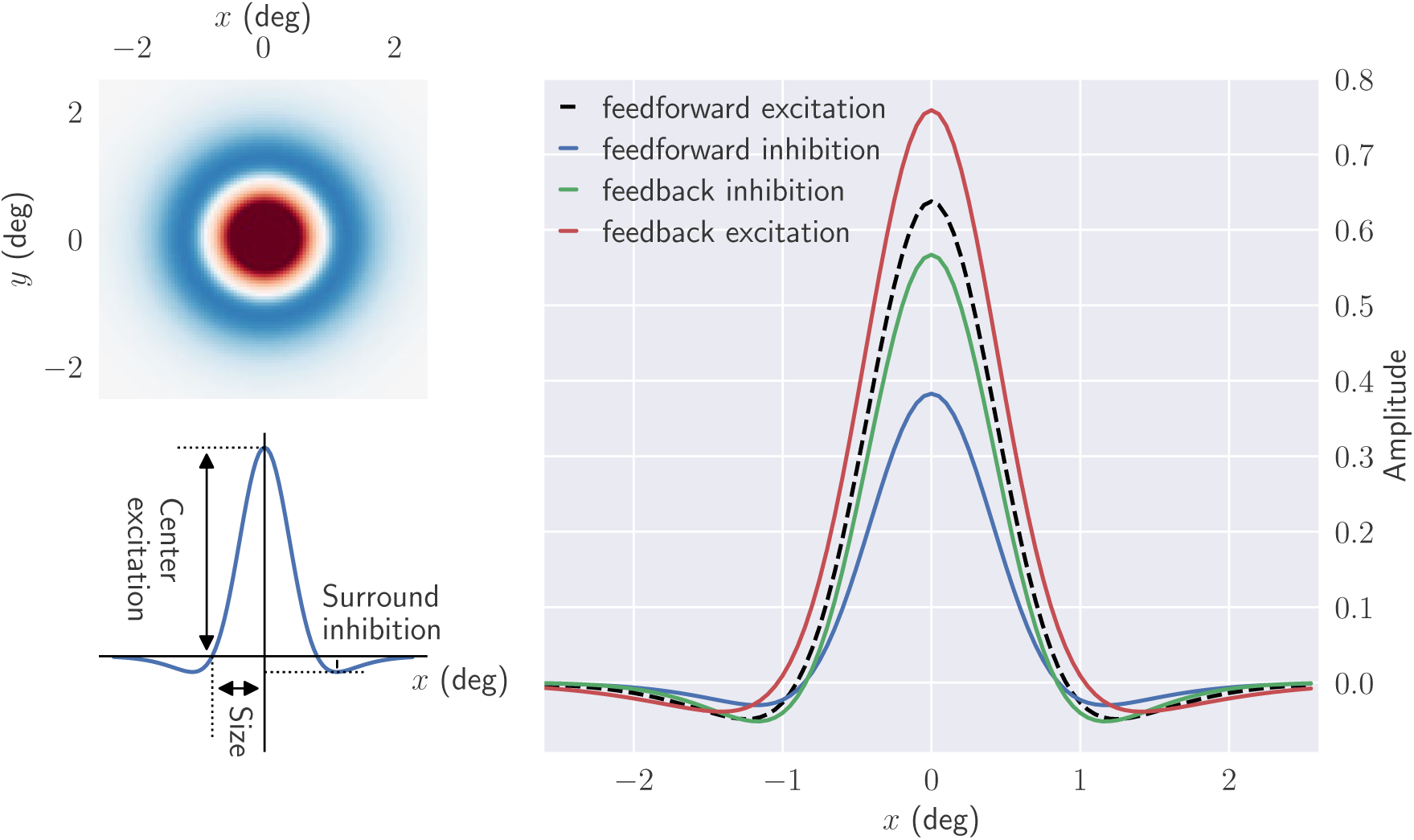
Cortical feedback modulates the center-surround receptive fields of relay cells. *Upper left*: the two dimensional spatial structure of the impulse-response function. *Bottom left*: one-dimensional plot of the impulse-response function. Center excitation and surround inhibition correspond to the maximum and minimum value of 
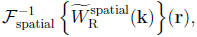
 respectively, while where the zero-crossing occurs is used as an indication for the receptive field size. *Right*: spatial impulse-response function for different circuit con. In each case all other contributions are removed, except feedforward excitation. Default parameters from Table 1 have been used.

Next, we investigate the spatial impulse-response function in more detail. In particular we study how the spatial responses depend on the weights (*w*) and Gaussian width parameters (*a*) of the connections, see left panel of Fig. 6. We characterize the spatial receptive-field structure by three measures: the receptive field size (radius), center excitation, and surround inhibition, cf. left panel of Fig. 5.

**Fig 6.**
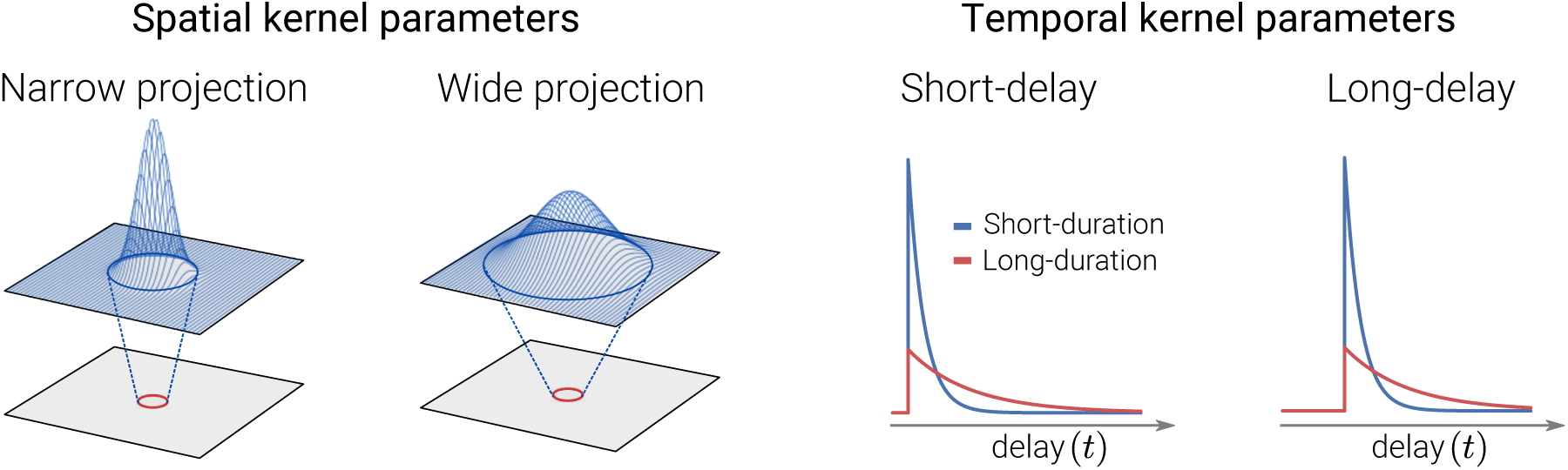
Illustration of spatial and temporal features of coupling kernels. *Left*: Spatial connectivity patterns between presynaptic neurons in the top layer and a single postsynaptic neuron (red circle) in the bottom layer for Gaussian width parameters (*a*). The Gaussian curves superimposed on top layers illustrate the spatial extent of the input to the neuron in the bottom layer. *Right*: Different scenarios for the temporal connectivity pattern. The time constant τ in the exponential decay function describes the duration, while Δ is the delay parameter. In the present example applications we have kept the time constant τ fixed at 5 ms.

In Figs. 7 and 8 the effect of kernel parameters on the spatial impulse-response function is shown for different circuit configurations. The effects of increasing feedforward inhibitory weight *w*_RIG_ and width *a*_RIG_ are shown in the top row of Fig. 7. The clear tendency is that narrow kernels with high weights most effectively reduce the center excitation and surround inhibition. The largest reduction in the receptive-field size is also observed in this situation. Another observation is that inhibitory kernels widths *a*_RIG_ similar to the width *b*_G_ ~ 1.3 deg of the DOG surround of the ganglion-cell input, combined with large weights, give a large surround inhibition.

**Fig 7.**
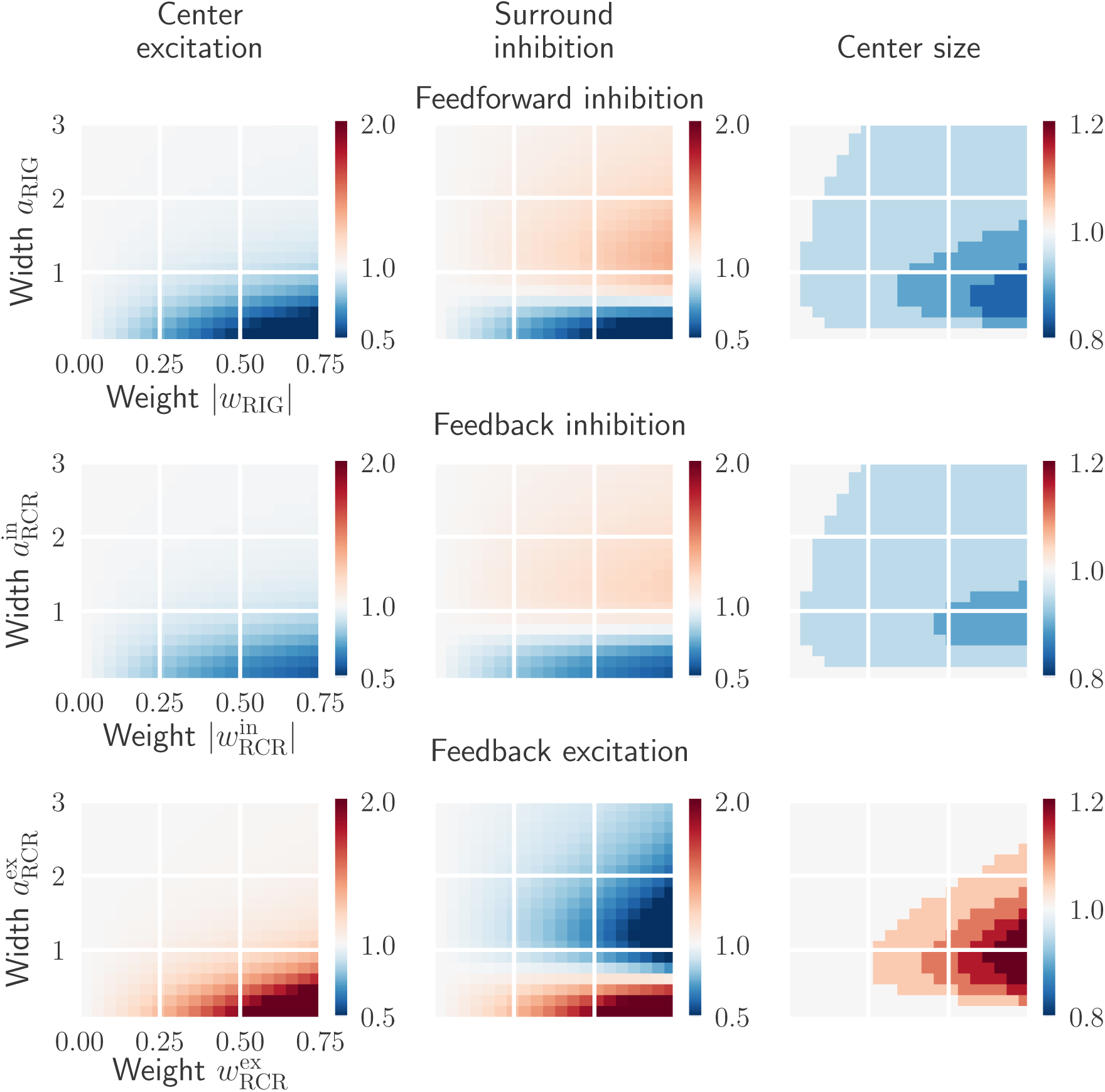
Effects on relay-cell spatial impulse-response function characteristics from excitatory and inhibitory inputs are opposite. *Top row*: dependence on the feedforward inhibition weight *w*_RIG_ and width *a*_RIG_. *Middle row*: dependence on the feedback inhibition weight 
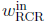
 and width 
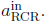
 *Bottom row*: dependence on the feedback excitation weigh 
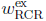
 and width 
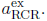
 All values are normalized with respect to the case where relay cells only receive feedforward excitation from retinal ganglion cells. The parameters in *W*_G_ and *K*_RG_ are kept fixed (see Table 1).

**Fig 8.**
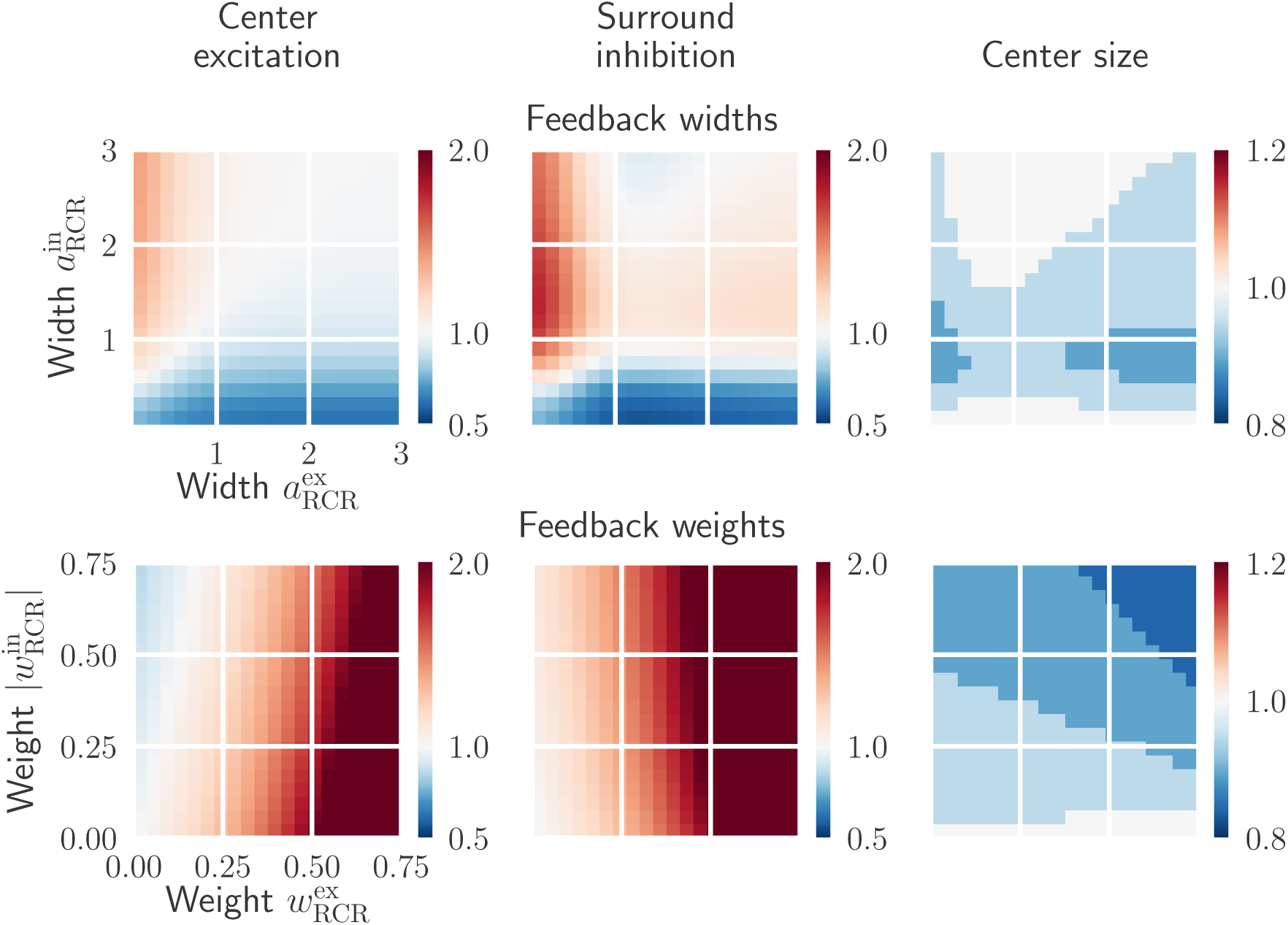
Effects on relay-cell spatial impulse-response function from mixed excitatory and inhibitory feedback. *Top row*: dependence on cortical feedback widths 
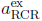
 and 
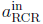
 with weights kept fixed: 
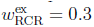
 and 
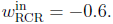
 *Bottom row*: dependence on cortical feedback weights *w*_RCR_^ex^ and 
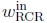
 with widths kept fixed: 
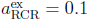
, and 
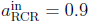
. All values are normalized with respect to the case where relay cells only receive feedforward excitation from retinal ganglion cells. The parameters in *W*_G_ and *K*_RG_ are kept fixed (see Table 1).

For the situation with feedback inhibition only (Fig. 7, middle row) an overall similar tendency is observed. However, the effects of feedback inhibition are somewhat weaker compared to feedforward inhibition for these example parameter ranges. Finally, we see from bottom row in Fig. 7 that strong and narrow excitatory feedback strongly increases the center excitation and surround inhibition. Larger widths, however, reduce the surround inhibition significantly and also results in larger receptive-field sizes.

To see the influence of a mixed cortical feedback on the spatial receptive-field properties, we show in Fig. 8 the effects of increasing cortical feedback weights and widths. A configuration consisting of a narrow excitatory and a broader inhibitory feedback both increases the excitation in the center and the inhibition in the surround. A large reduction in the receptive-field center size is also seen with this configuration, specially for inhibitory width values 
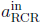
 close to one.

The bottom row of Fig. 8 shows the effect of increasing feedback weights for a narrow excitatory central core projection and a wider inhibitory projection. Strong excitatory feedback combined with a weak inhibitory feedback increases excitation in the center and inhibition in the surround. In contrast, strong inhibitory feedback combined with weak excitatory feedback, reduces the center excitation. Note however that the effects due to strong excitatory feedback are more significant than the ones due to the inhibitory feedback. This is specially obvious in the surround inhibition which is nearly completely dominated by the excitatory feedback strength. The size of the receptive field decreases with increasing excitatory and inhibitory feedback strength.

In conclusion, these results show that the cortical feedback is well suited to modulate the center-surround organization of relay-cell receptive fields. Excitatory and inhibitory inputs have opposite effects on a relay cell’s spatial response: while excitatory feedback can increase the center excitation and center size, inhibitory feedback can do the opposite. Depending on the width of the feedback projection, both excitatory and inhibitory feedback can either increase or decrease surround excitation. A mixed feedback configuration consisting of a combination of narrow excitatory and a broader inhibitory feedback, both increases the excitation in the center and the inhibition in the surround.

#### 3.1.2 Area summation curves

A common way to experimentally probe the center-surround organization of cells in the early visual pathway is to measure *area-response curves*, i.e., the response to circular stimulus spots as a function of spot diameter [16, 17, 22, 56, 60, 79–81]. In Fig. 9 we correspondingly show area-response curves for relay cells responding to static bright-spot stimuli for different feedback configurations. Here the receptive field of the cell is set to be concentric with the spot.

**Fig 9.**
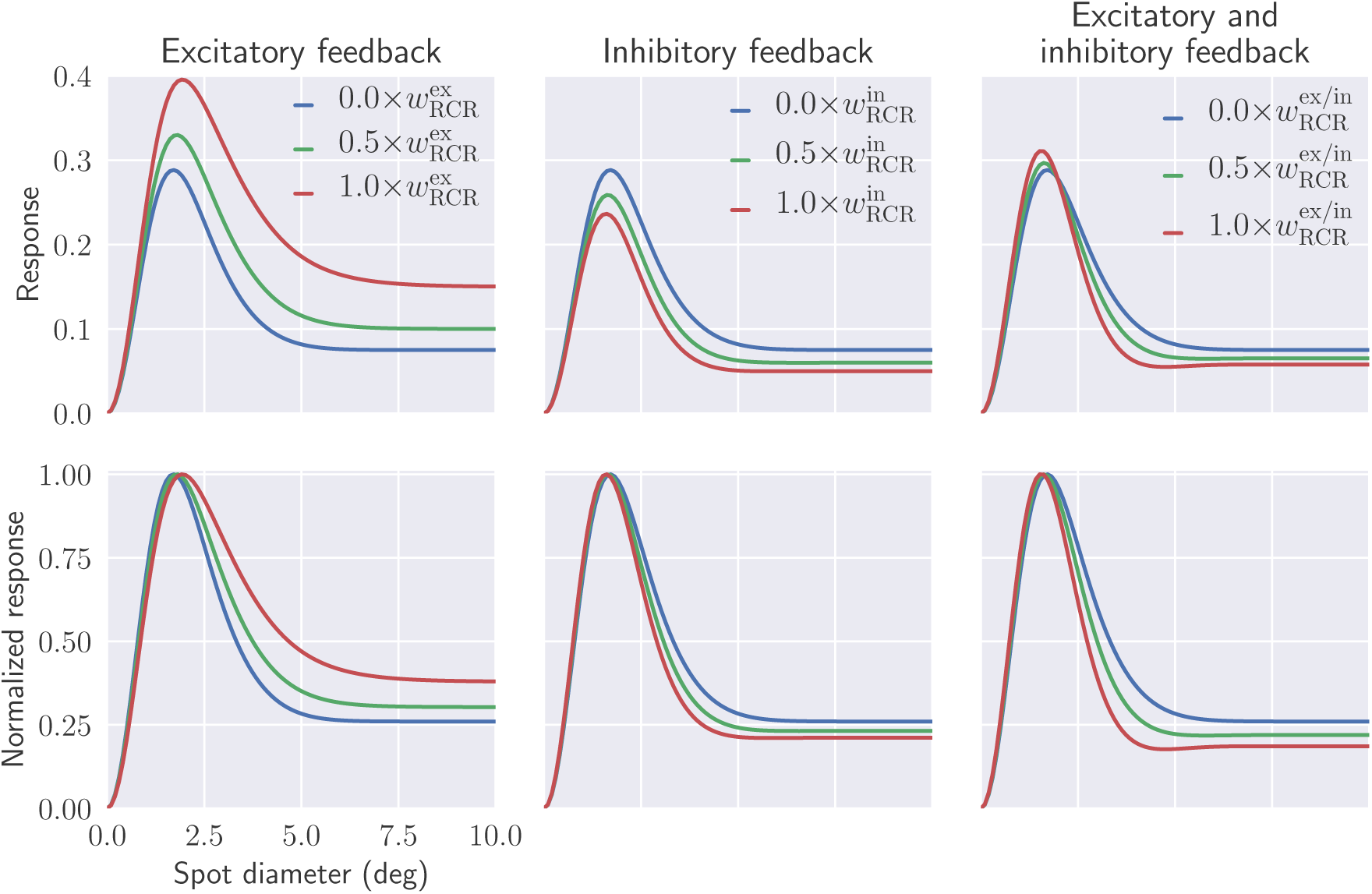
Mixed feedback may enhance both excitatory response to stimuli within the receptive-field center (unlike inhibitory feedback alone), and suppressive effects of stimuli in the surround (unlike excitatory feedback alone). Predicted area-response curves of relay cells for different arrangements of cortical feedback. Bottom row shows normalized curves. Default values from Table 1 have been used for fixed parameters.

The first column in Fig. 9 shows that an increasing excitatory feedback enhance the excitatory response to stimuli restricted to be within the receptive field center. It also reduces the suppressive effects of stimuli in the surround area. Inhibitory cortical feedback, on the other hand, reduces the response to optimal patch diameter and enhances the suppressive effects for large patch sizes (second column in Fig. 9).

In the last column of Fig. 9, the mixed feedback situation with a combination of narrow excitatory and a broader inhibitory feedback, as suggested by experimental findings [3,16], is considered. Here we observe that an increased feedback strength both (i) enhances the excitatory response to stimuli restricted to be within the receptive field center, and (ii) enhances the suppressive effects of stimuli in the surround area. Stronger feedback also reduces the receptive-field center size, i.e., the spot diameter giving the maximum response.

The area-response curves in Fig. 9 are for static spot stimuli, but area-response curves are also commonly recorded for patch-grating stimuli [17,22,39,80]. In our formalism such response curves are readily obtained by use of the circular patch-grating stimulus function *S* in Eq. (20). The resulting area-response curves typically resemble the static-spot curves shown in Fig. 9, and we do not show any example curves here.

However, in Fig. 10 we summarize results for area-response curves both for static-spot (Fig. 9) and patch-grating stimuli. Here, the stimulus size giving the largest response (corresponding to the receptive-field center size for static spot stimuli) and center-surround suppression index are shown as a function of feedback strength. This suppression index *α*_s_ is here defined as:

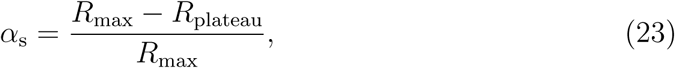

**Fig 10.**
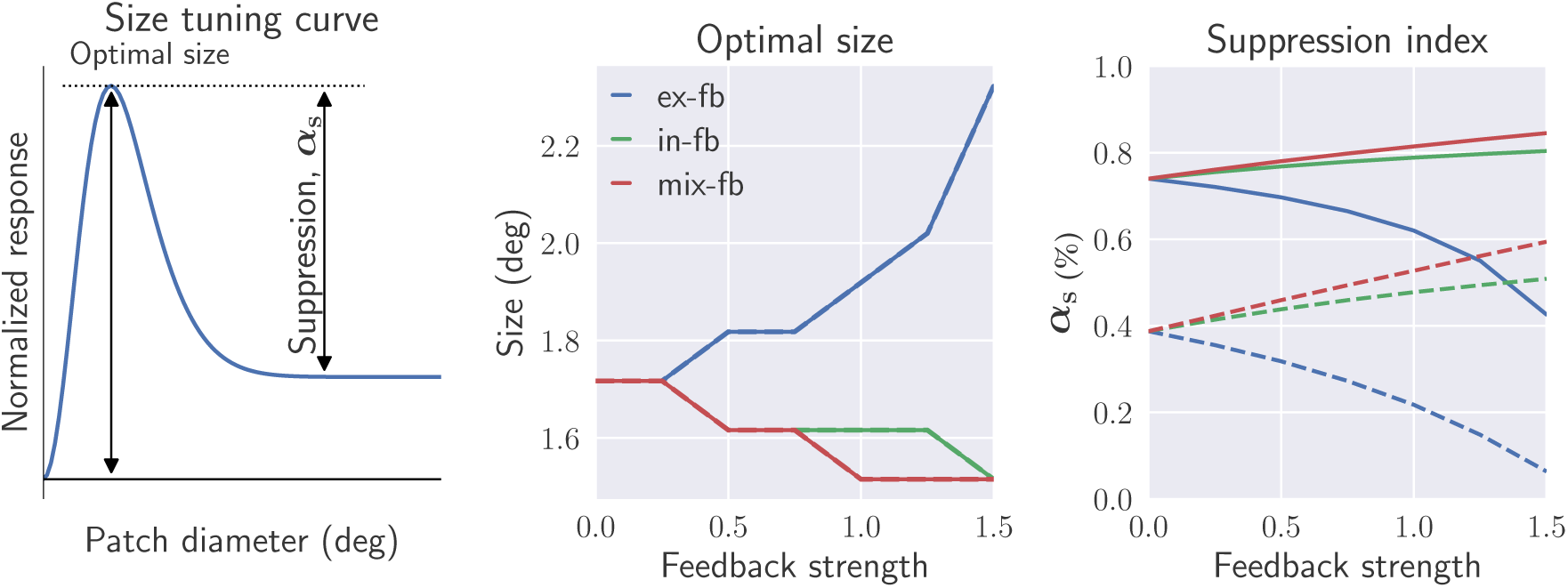
Optimal stimulus size and suppression index decrease and increase with increased inhibitory and (present) mixed feedback, respectively, while with excitatory feedback trends are opposite. Optimal size and suppression index (α_s_) are shown as a function of cortical feedback weight for different feedback configurations. These are extracted from the size tuning curve using static spot (solid lines) and patch grating (dashed lines, 
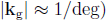
 as stimulus (left most figure). The values on the *x*-axis represent factors multiplied with the default values for 
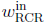
 and 
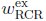
 listed in Table 1. Default values for fixed parameters are also listed in this table.

where *R*_max_ is the maximum response, and *R*_plateau_ is the response when the large-diameter plateau is reached (see left panel in Fig. 10).

The figure shows that the suppression index for static spot stimuli (right panel, solid lines) is increased with stronger feedback weights both for inhibitory and mixed feedback. The same qualitative trend is also observed for patch-grating stimuli (dashed line). Here the suppression index without feedback is fairly small (~0.4), but increases more strongly with feedback strength than for static-spot stimuli. This relative difference in suppression index between spot and patch-grating stimuli is qualitatively in agreement with experimental observations [17,22]. With excitatory feedback on the other hand, the suppression index is reduced with increasing feedback strength. The largest suppression indices are found for mixed feedback, again illustrating that such feedback is particularly suited for modulating center-surround antagonism.

The optimal stimulus size (Fig. 10, middle panel) is seen to be the same for static-spot and patch-grating stimuli. For both stimulus types this size is seen to decrease with increasing feedback strength both for inhibitory and mixed feedback, while the opposite is true for excitatory feedback.

Note that in experimental measurements of spot area-response curves, ‘flashing’ spots rather than static spots have been used [22, 60, 79]. This means that the spots were ‘flashed’ on and the subsequent response, which contained both a transient and a sustained response, were used to compute the area-response curves [79]. The static-spot area-response curves computed here would correspond to the sustained response, but the area-response curves based on the transient part or the sustained part are expected to have similar shapes, see Fig. 13 in [82]. For the patch-grating experiments in [17, 22] drifting patch gratings with a temporal frequency of only *ω*~6 Hz was used, so that the ‘fast-loop limit’ (i.e., assuming sufficiently short propagation times around the thalamocortical loop), the expression in Eq. (22) expectedly still can be used, see discussion in [40].

#### 3.1.3 Spatial frequency tuning curves

The spatial summation curves in Fig. 9 show that the cortical feedback modulates the size tuning properties of relay cells. Next, we investigate the influence of cortical feedback on spatial frequency tuning of relay cells. In Fig. 11 the tuning curves at two different patch sizes are shown. The smaller patch is similar in size to the receptive-field center size, while the larger patch covers both the center and surround of the receptive field.

**Fig 11.**
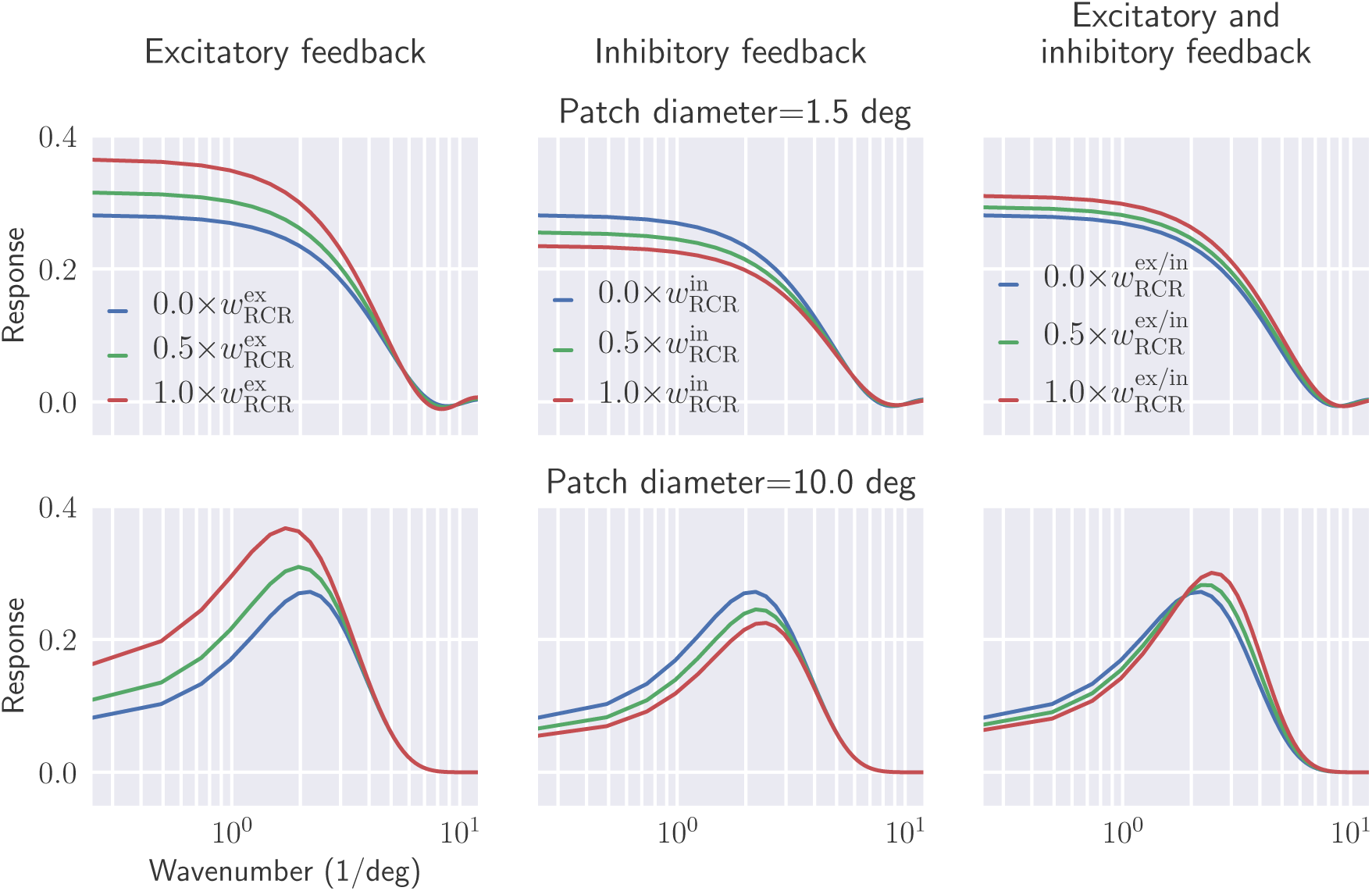
Shift from low-pass to band-pass characteristics is seen in spatial frequency tuning of relay cells when increasing stimulus patch size. Wavenumber 
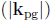
 tuning of relay cells, using patch grating at two different patch sizes (rows), is shown for different feedback configurations (columns). Default values from Table 1 have been used for fixed parameters.

For the smaller patch the frequency characteristic corresponds to a low-pass filter for all feedback configurations, cf. upper row panels in Fig. 11. Increasing feedback strength leads as expected to higher response values for excitatory feedback, but also for the mixed feedback. For inhibitory feedback the opposite is the case.

For the larger patch size, (effectively corresponding to a full-field grating), relay cells have band-pass characteristics in all cases, cf. lower row panels in Fig. 11. Excitatory feedback is seen to overall increase the response as well as shift the frequency giving the maximum to smaller frequencies. Inhibitory feedback is seen to have opposite effects. For mixed feedback an interesting combination of these effects are seen, i.e., the maximum-frequency response is shifted towards higher frequencies, but the maximum amplitude is also increased.

The shift from low-pass to bandpass characteristics when changing the grating size can be explained by considering the center-surround organization of the receptive field. When the stimulus only covers the center of the relay-cell receptive field, the filtering of the circuit is effectively Gaussian-like, i.e., a low-pass filter. However when the stimulus also covers the surround region, the circuit filter is effectively an antagonistic center-surround filter with bandpass characteristic.

A putative benefit of the shift of the response towards higher frequencies observed for our mixed feedback, can be alluded to in the context of information theory and efficient coding. In natural scenes there are usually extensive spatial correlations since neighbouring regions often have similar luminance values [29]. This leads to a power spectrum of the input with large contributions from low spatial frequencies. An antagonistic center-surround organization dampens the low-frequency components and enhances the higher frequency components of the image and reduces the redundancy in the signal conveyed to the cortex. The shift towards higher frequencies sharpens the spatial receptive field of the relay cells and thereby increases the saliency of edges. To illustrate this point, we show in Fig. 12 the response map for the relay cells for different circuit configurations with a natural image as stimulus. It is clear that in this example the mixed cortical feedback improves the detection of edges considerably.

**Fig 12.**
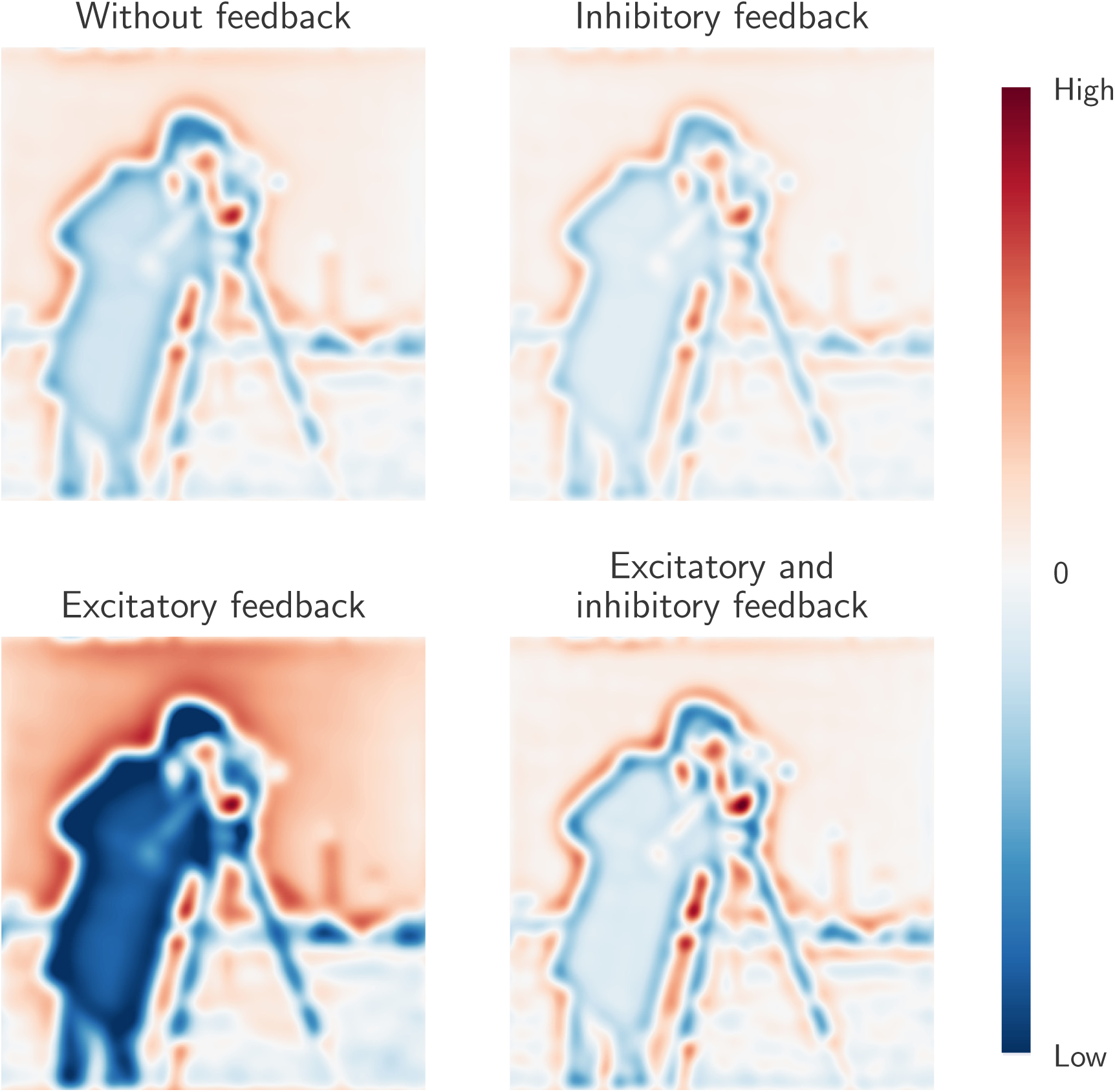
Mixed feedback has different effect on low and high frequency components of natural scenes in contrast to pure excitatory or inhibitory feedback. Each subfigure shows activation of a layer of relay cells in response to the input image, shown as a heatmap from blue to red (low to high response). Default values from Table 1 have been used except for the feedback weights 
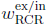
 which have been set at 1.5 times the listed default values.

### 3.2 Effect of cortical feedback on temporal properties

#### 3.2.1 Temporal receptive fields

We have so far focused on influence of cortical feedback on spatial response properties of relay cells. Next, we investigate the effect of cortical feedback on temporal properties. The relay cell impulse-response function in Fourier space in Eq. (14) can be written as

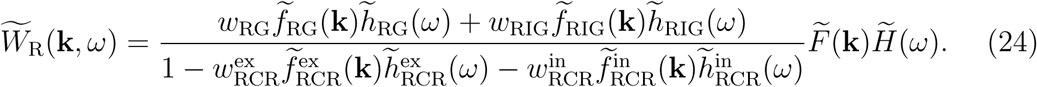

Here, 
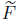
 and 
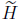
 represent the assumed spatial and temporal response functions for the retinal input, i.e., Eq. (15) and Eq. (16), respectively. Further the coupling kernels have been expanded into products of spatial 
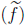
 and temporal 
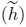
 functions, cf. Eq. (17).

Illustrations of the real-space version *W*_R_(**r**, *t*) of this impulse-response function is shown in Fig. 13 using the kernel parameters listed as default parameters in Table 1. In this figure the temporal evolution of the spatial structure of the receptive field is shown (top panel), in addition to an *x*-t plot of the impulse-response function, summarizing how the one-dimensional spatial organization of the receptive field changes with time. These figures illustrate the biphasic nature of the center and surround responses, as has been observed experimentally [62,83]. For *t* between 0 and 50 ms the impulse-response function exhibits a bright-excitatory center, i.e., an increased firing to a tiny bright test spot placed in the receptive-field center. However, for times later than 60 ms, the polarity of the center response is reversed, and becomes dark-excitatory, i.e, increased firing-rate for dark spots. A similar behavior is observed for the surround, but with opposite polarities.

**Fig 13.**
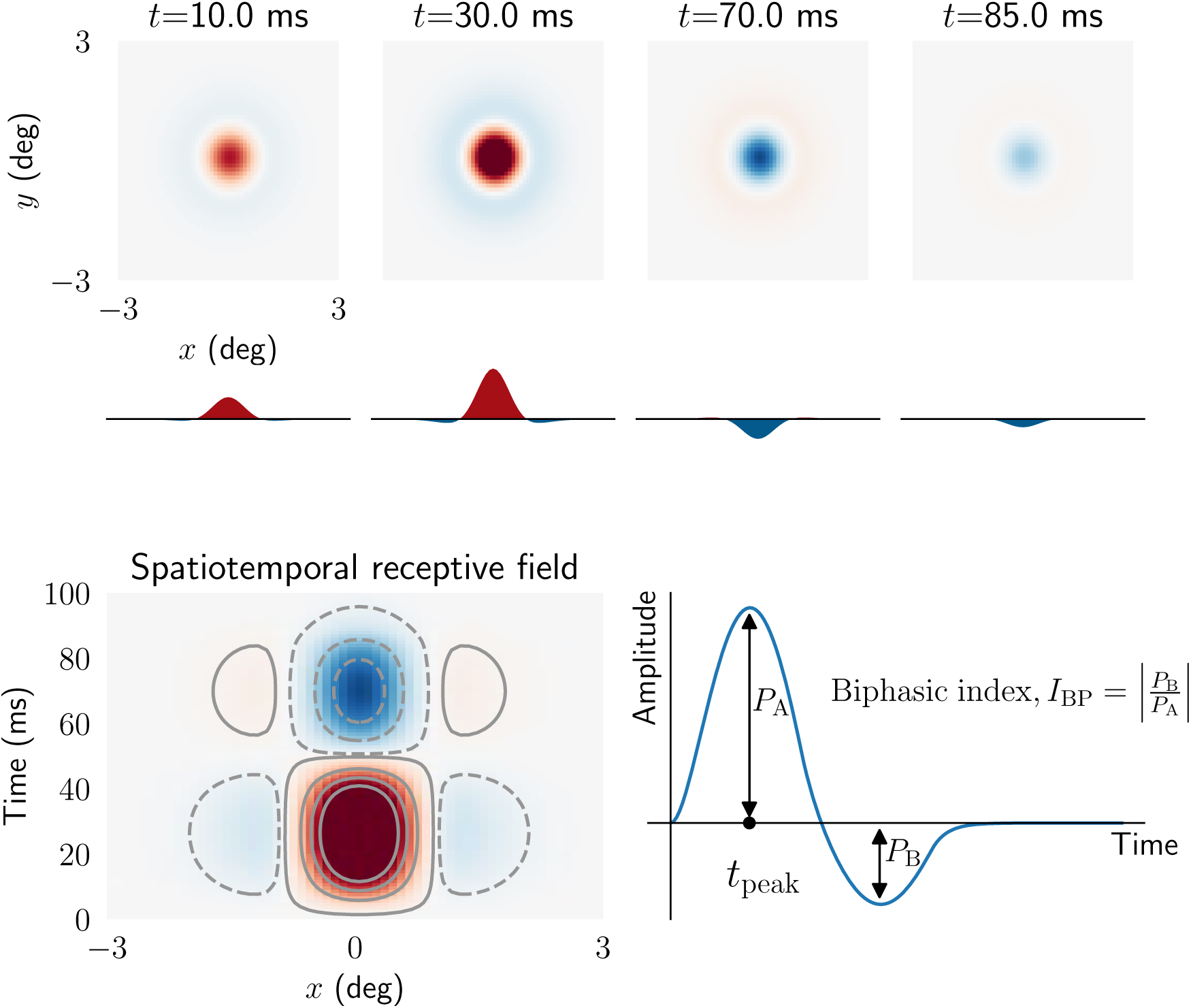
Spatiotemporal impulse-response function of relay cell. *Top*: panels showing spatial receptive field at different times. Curves below panels are one-dimensional plots of the receptive fields as a function of *x* alone. *Bottom left: x-t* plot of receptive field. ON regions are shown in red (solid lines) while OFF regions are shown in blue (dashed lines). *Bottom right*: Curve showing temporal evolution of ON region of the receptive field. The biphasic index *I*_BP_ is defined as the ratio between the peak magnitude of the (negative) rebound phase and the peak magnitude of the first (positive) phase. *t*_peak_ is the peak response latency. Note that only feedforward excitation from retinal ganglion cells to relay cells is included.

Experimental studies have reported several distinct temporal receptive-field profiles among relay cells, including monophasic and triphasic responses in addition to the more common biphasic response seen in Fig. 13 [72, 84]. To explore scenarios where these different response profiles may arise, we next study how different model parameters change the shape of the real-space temporal impulse response. In accordance with previous experimental and computational studies we use the *biphasic index* (*I*_BP_) and p*eak response latency* (*t*_peak_) as measures to characterize the temporal properties of the impulse-response function [54,62,84]. The biphasic index is defined as the ratio between the peak magnitude of the (negative) rebound phase and the peak magnitude of the first (positive) phase, and thus measures how biphasic the response is (Fig. 13, bottom right). A biphasic index equal to one means a perfect biphasic response, while zero corresponds to a monophasic response.

Fig. 14 illustrates the dependency of the temporal part of the impulse-response function, as well as *I*_BP_ and *t*_peak_, on various circuit configurations and model parameters. Each temporal coupling kernel is described by two parameters: the time constant *τ* of the exponential decay and the parameter Δ accounting for delay in the propagation of the signals between the different neuronal population. In the present examples we keep the *τ* fixed and instead focus on how different values of the Δ affect the temporal impulse-response function. In Fig. 14 it is seen that the effects of feedforward and feedback inhibition on the temporal impulse are qualitatively similar. In both cases the depth of the second phase is increased for delayed (large Δ) inhibitory input, i.e., the biphasic index *I*_BP_ is increased. The peak response latency *t*_peak_ is seen to be substantially reduced with feedforward inhibition, but less so for feedback inhibition. Finally, we see from the middle left panels in Fig. 14 that for strong delayed inhibitory inputs, a triphasic impulse-response may arise.

**Fig 14.**
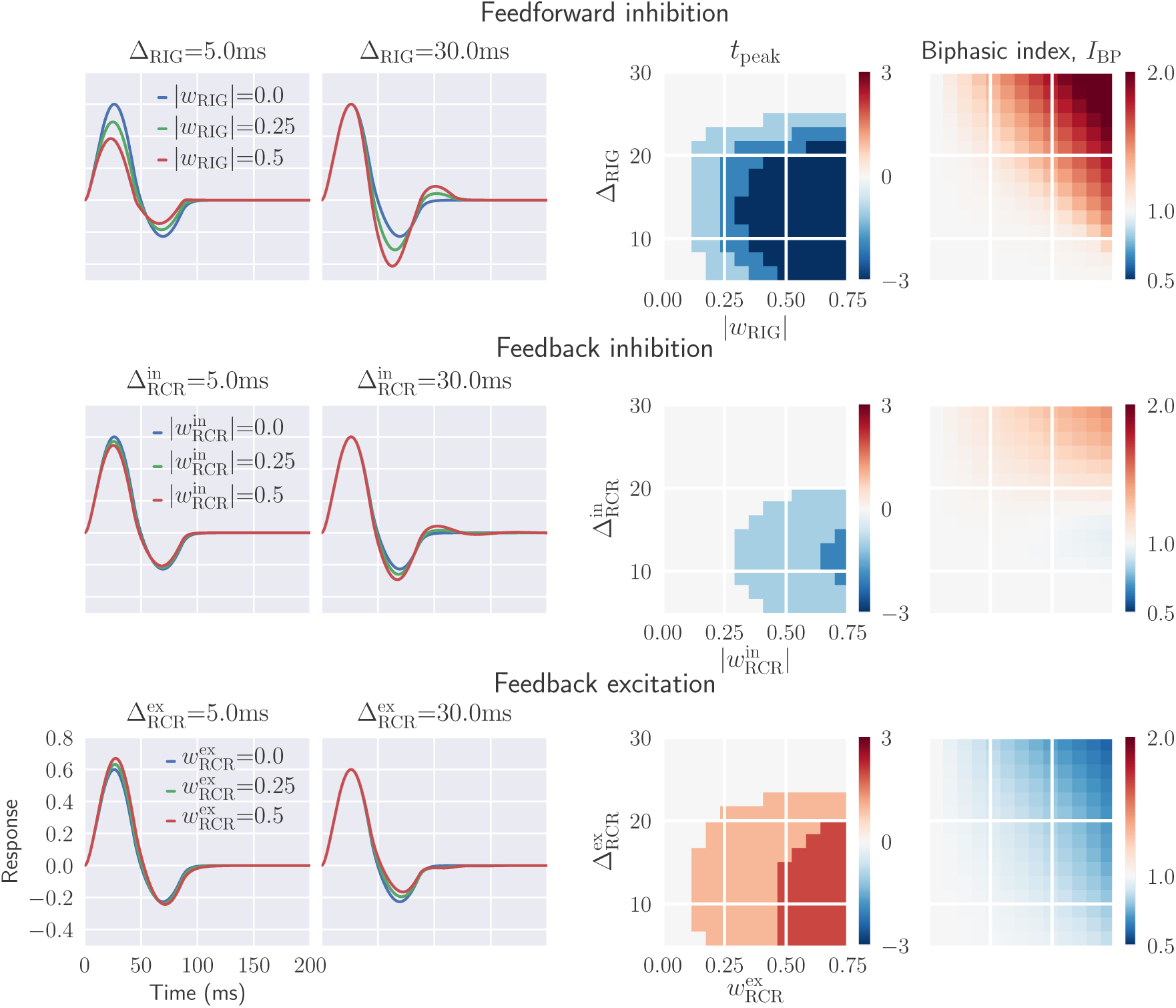
Inhibitory feedback can increase the biphasic index and reduce the peak response latency, while the opposite is seen for excitatory feedback. *Left panels*: temporal evolution of the relay-cell impulse-response function (ON region) for different circuit configurations, i.e., feedforward inhibition only (top), feedback inhibition only (middle), feedback excitation only (bottom). The feedforward excitation is fixed in all cases. *Right panel*: parameter dependence of two impulse-response measures *t*_peak_ and biphasic index *I*_BP_. The biphasic index is normalized with respect to the value for the case with feedforward excitation only (*I*_BP_ = 0:35), while the *t*_peak_ plots show the difference in peak time in milliseconds compared to the corresponding value for feedforward excitation only (*t*_peak_ = 29 ms). Default parameters have been used for the fixed parameters (see Table 1)

Focusing on the middle and bottom row of panels in Fig. 14, we see that the effect of excitatory feedback is essentially the opposite of that for inhibitory feedback. The biphasic index *I*_BP_ is decreased by the excitatory feedback and increased by the inhibitory feedback, in particular for delayed inputs. Further, the peak response latency t_peak_ is increased with excitatory feedback, in contrast to with inhibitory feedback.

In Fig. 15 the effect of temporal kernel parameters on the biphasic index *I*_BP_ and *t*_peak_ is shown for the more complex situation with a mixed excitatory and inhibitory feedback. Here the spatial spread as well as the relative weight of the excitatory and inhibitory feedback is kept fixed while the delay parameters 
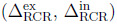
 are varied. With long-delay excitatory feedback and short-delay inhibitory feedback 
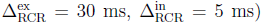
, the first positive phase of the impulse-response function is only modestly affected by the feedback. In this case the feedback mainly affects the second negative phase, which generally is reduced in depth. Thus the response becomes more monophasic, as reflected in smaller values for the biphasic index (Fig. 15, rightmost panel).

**Fig 15.**
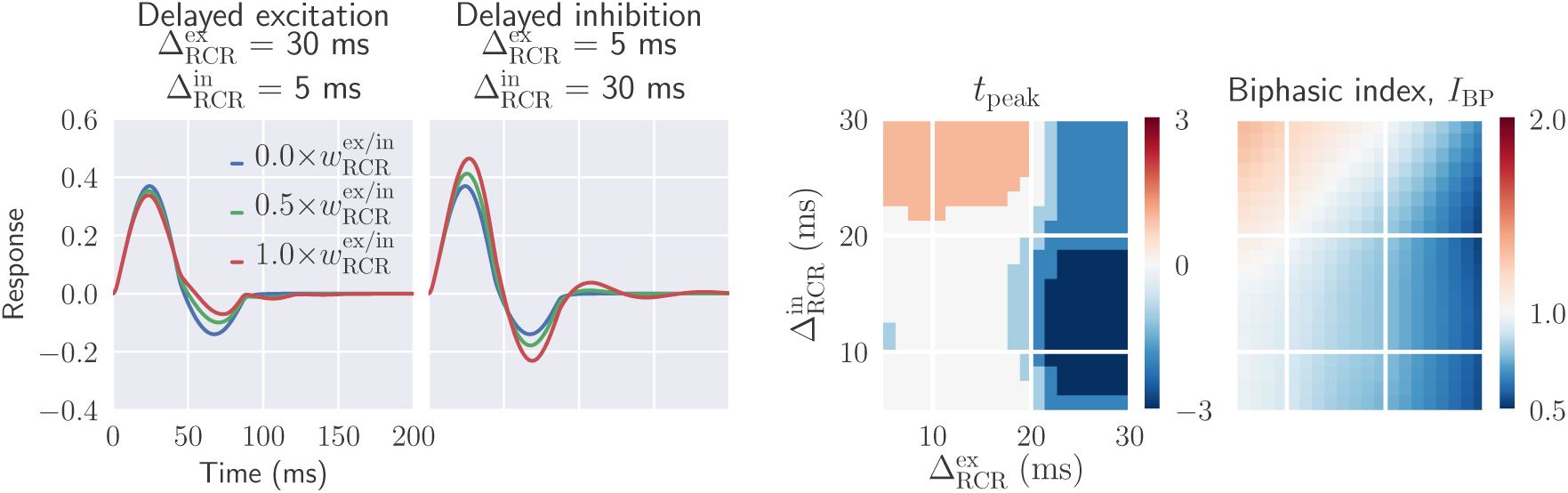
Mixed feedback: delayed inhibitory feedback gives oscillatory responses, delayed excitatory feedback more monophasic responses. *Two leftmost panels*: temporal impulse-response function with mixed excitatory and inhibitory feedback, where feedforward inhibition also is included. *Two rightmost panels*: parameter dependence of two impulse-response measures *t*_peak_ and biphasic index *I*_BP_. See Fig. 14 caption for details. Default parameters have been used for fixed parameters (see Table 1)

In the case with long-delay inhibitory feedback and short-delay excitatory feedback 
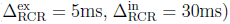
, the rapid excitatory feedback boosts the initial positive peak while the former boosts the following negative peak. Further, this feedback combination gives multiphasic responses, i.e., distinct responses also after the initial biphasic response.

The figures in the right panel of Fig. 15 show that the peak response latency *t*_peak_ and the biphasic index *I*_BP_ are reduced for long-delayed excitatory feedback. Long-delayed inhibitory feedback combined with short-delayed excitatory input, on the other hand, increases both *t*_peak_ and *I*_BP_.

#### 3.2.2 Temporal frequency tuning curves

To investigate the effect of cortical feedback further, we next explore the temporal frequency tuning of relay cells. This is done by computing the response of the relay cells to a full-field grating stimulus for a range of different temporal frequencies [85–88]. In this case the response is given by the magnitude of the Fourier-space impulse-response function in Eq. (24) evaluated with a varying angular frequency *ω*_g_ combined with a fixed spatial wave vector **k**_g_ [76]. In the present example the wavenumber is kept fixed at 
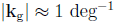
.

In Fig. 16 the frequency tuning is shown for inhibitorycitatory feedback (bottom). As expected, rapid inhibitory feedback 
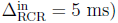
reduces the overall response. In addition, it is also seen to shift the peak frequency to slightly higher values. Long-delayed inhibitory feedback 
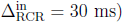
, on the other hand, gives sharper tuning curves, enhancing the band-pass characteristics of relay cells. In this case increased feedback weights both sharpen the resonance and shift it to higher frequencies.

**Fig 16.**
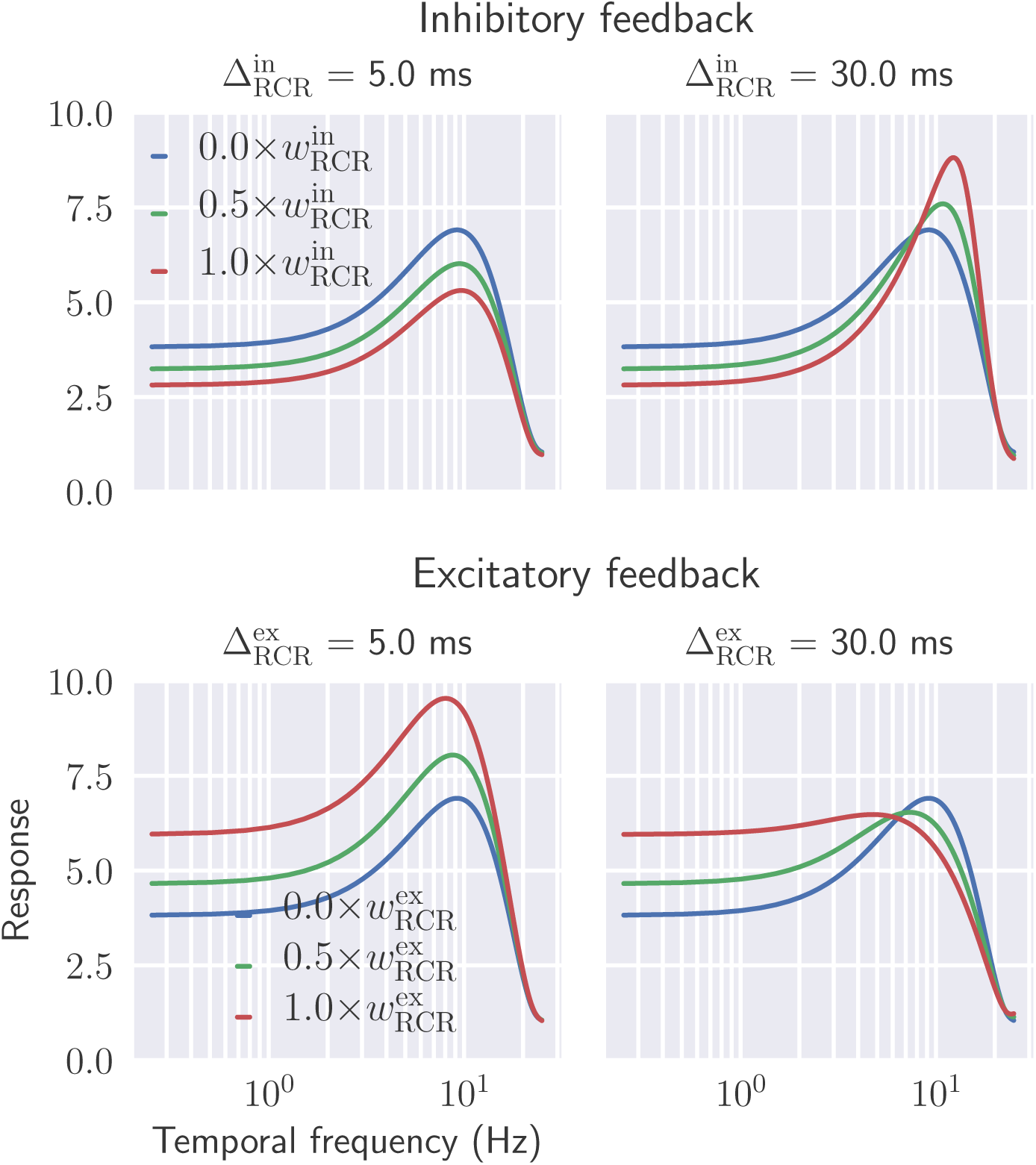
Delayed inhibitory feedback sharpens the temporal frequency tuning of relay cells, while delayed excitatory feedback blunts the temporal frequency tuning of relay cells. Effect of cortical feedback on temporal frequency tuning of relay cells are shown for different values of thalamocortical delay. *Upper row*: Inhibitory feedback only. *Lower row*: Excitatory feedback only. Full-field grating is used as stimulus 
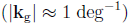
 and default values from Table 1 is used for fixed parameters.

Excitatory and inhibitory feedback essentially have opposite effects on the tuning properties of relay cells (Fig. 16, bottom row). Rapid excitatory feedback shifts the tuning curve to higher response values, while long-delayed excitatory feedback blunts the tuning profile. In all cases the peak frequency is shifted to lower frequencies.

In Fig. 17 we next investigate the effect of mixed cortical feedback on temporal frequency tuning properties of relay cells. In particular, we consider three different cases: (1) long-delay inhibitory feedback combined with short-delay excitatory feedback, (2) long-delay excitatory feedback combined with short-delay inhibitory feedback, and (3) synchronized feedback where excitatory and inhibitory feedback are received at the same time. Long-delayed inhibition (combined with rapid excitatory feedback) leads to sharper tuning with increasing feedback. This contrasts the case with long-delay excitation (combined with rapid inhibitory feedback), where a more flat spectrum is observed with increasing feedback. Synchronized excitatory and inhibitory feedback does not change the tuning properties by much when comparing with the case without cortical feedback. This reflects that in this case the excitatory and inhibitory effects largely cancel each other. Note that this cancellation is not perfect since the weight and spatial properties of the two feedback types are not identical, cf. Table 1.

**Fig 17.**
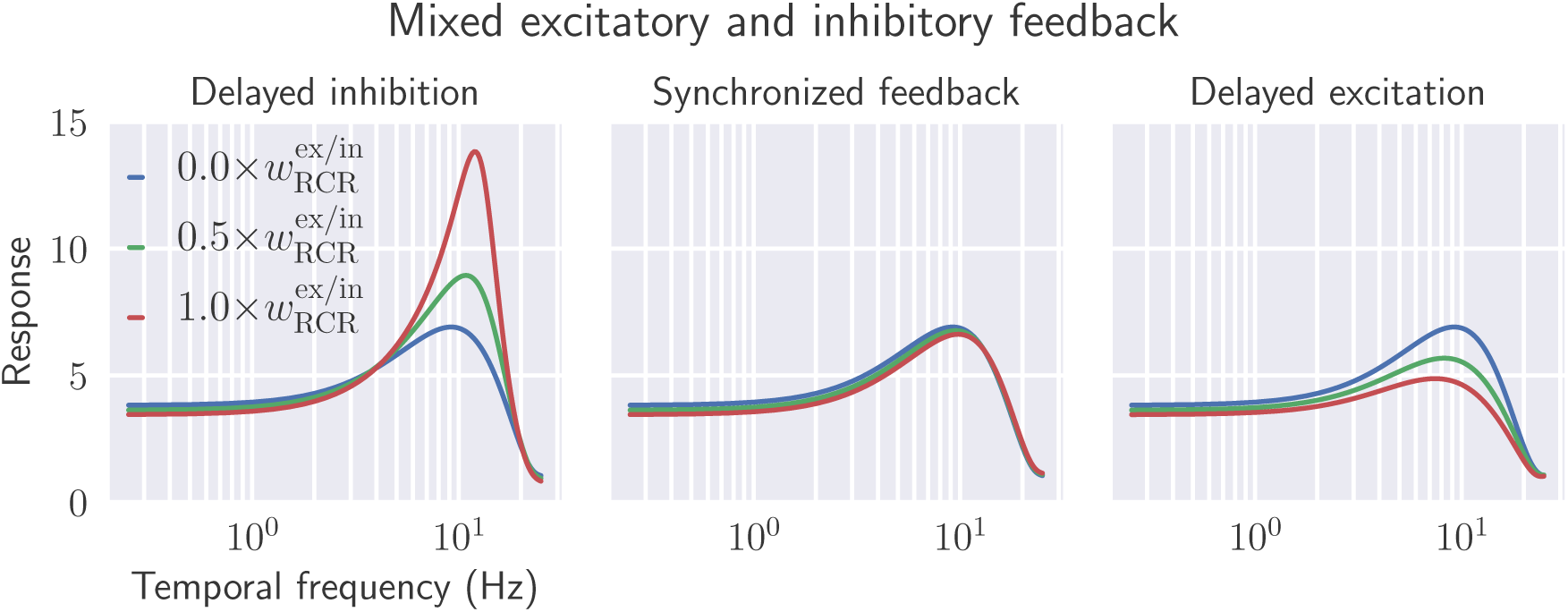
Mixed feedback: delayed inhibitory feedback relative to excitatory feedback sharpens the temporal frequency tuning of relay cells, while the opposite blunts the tuning. *Left*: Delayed inhibition, i.e., rapid excitatory feedback combined with long-delay inhibitory feedback 
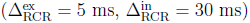
. *Middle*: Synchronized feedback, i.e., excitatory and inhibitory feedback arrive simultaneously 
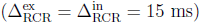
. *Right*: Delayed excitation, i.e., long-delay excitatory feedback combined with rapid inhibitory feedback 
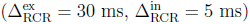
. Full-field grating is used as stimulus 
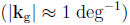
 and default values from Table 1 are used for fixed parameters.

In conclusion, the tuning curves from Figs. 16 and 17 show that both temporal low-pass filtering and band-pass filtering can arise from cortical feedback. The detailed spectral shape will depend on both the relative weight and relative delay of the excitatory and inhibitory feedback contributions. While long-delay inhibitory feedback sharpens the temporal frequency tuning of relay cells giving more band-pass-like characteristics, long-delay excitatory feedback makes the tuning more low-pass like. These tuning behaviors can be related to the temporal impulse-response functions depticted in Figs. 14 and 15: multiple phases in the temporal response leads to band-pass filtering, while monophasic responses have low-pass filter characteristics.

Note, finally, that the results in Figs. 14 to 17 are all obtained with temporal kernels with a fixed, relatively short, time constant *τ* of 5 ms, i.e., a relatively short duration of the feedback, cf. Fig. 6. With a longer duration, i.e., larger value of *τ*, qualitatively similar results for both the impulse-response function and temporal frequency tuning are found, though the curves were found to be more blunt.

#### 3.2.3 Decorrelation of naturalistic stimuli

In natural visual scenes there are, in addition to extensive spatial correlations, also large inter-frame correlations [42,89]. This means that pixels have a luminosity which usually changes gradually in time. It has previously been shown that the biphasic temporal response, seen in retina and dLGN, decorrelates the incoming signal in time, resulting in a more efficient representation of the information in the natural-scene images [42,90–92]. It has also been suggested that the cortical feedback may control the degree of temporal decorrelation in relay cells depending on the signal to noise ratio [37,42].

Here we investigate the putative role of cortical feedback on the decorrelation of visual input by calculating the temporal autocorrelation function for relay cell responses to natural movies for different feedback arrangements. The movie was recorded by a camera mounted on the head of a cat exploring the environment [93,94]. The average stimulus autocorrelation function in time and the averaged response autocorrelation function for the relay cells, is shown in Fig. 18.

**Fig 18.**
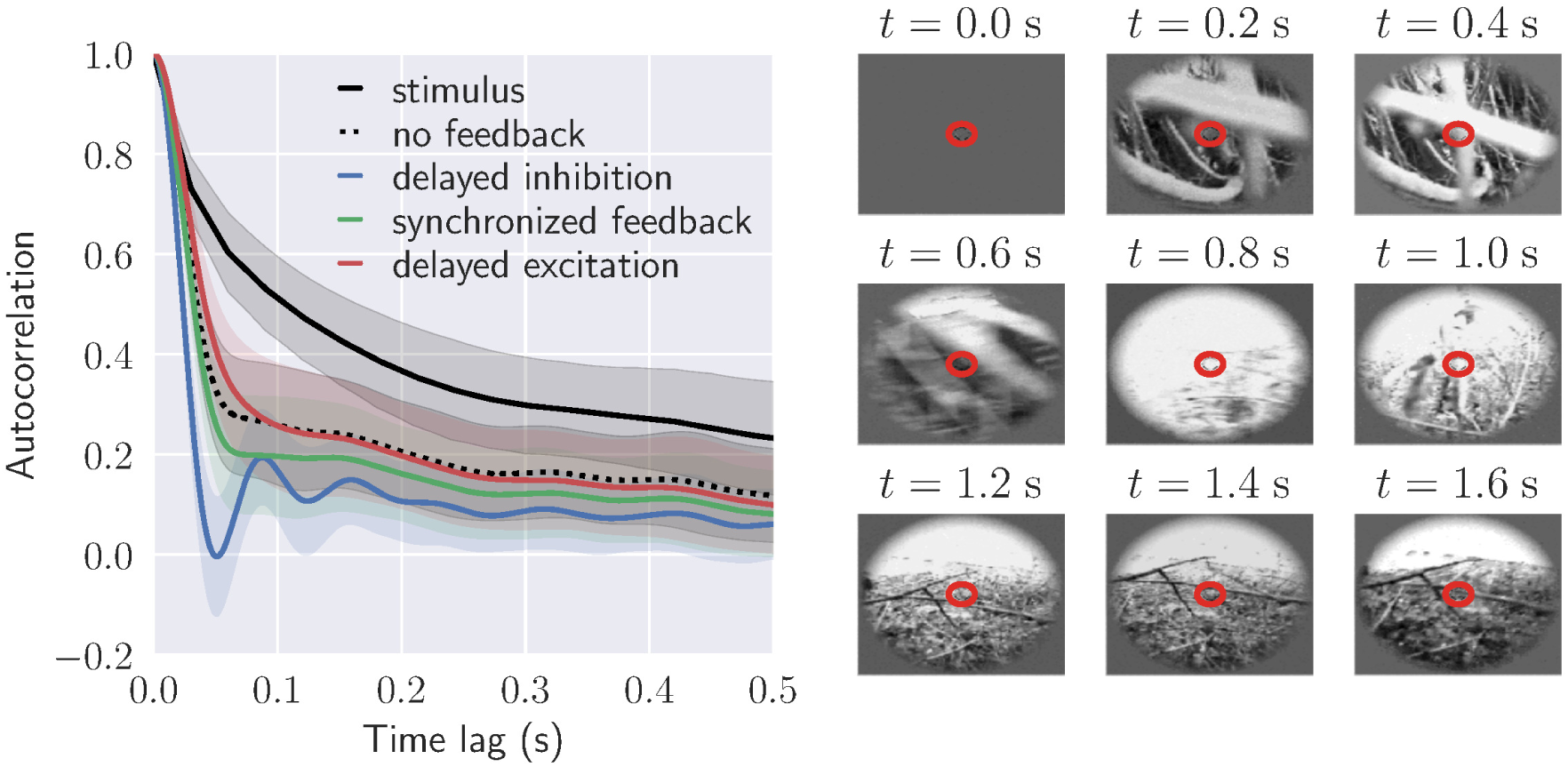
Cortical feedback may control the degree of temporal decorrelation in relay cells. *Left*: Autocorrelation function of stimulus and relay cell response for different circuit configurations: no feedback, long-delay inhibitory feedback combined with short-delay excitatory feedback, long-delay excitatory feedback combined with short-delay inhibitory feedback, synchronized feedback. In each case the average autocorrelation from 40 × 40 neurons at the center is shown with corresponding standard deviation. Default values from Table 1 have been used for fixed parameters, and the temporal feedback parameters are the same as in Fig. 17. *Right*: Frames from the complex naturalistic movie used as stimulus. This movie was recorded by a camera mounted on the head of a cat exploring the environment (forest) [93,94]. The red circle marks the receptive-field center size for the relay cell at the center.

The correlation has been calculated for both cases with and without cortical feedback. For the case with cortical feedback the three mixed-feedback scenarios from Fig. 17 are considered: (1) long-delay inhibitory feedback combined with short-delay excitatory feedback, (2) long-delay excitatory feedback combined with short-delay inhibitory feedback, and (3) synchronized feedback where excitatory and inhibitory feedback are received at the same time.

Fig. 18 shows that even without cortical feedback, the correlations in the relay-cell response are significantly lower than the correlations in the stimulus. We observe that the response correlations are further reduced by synchronized cortical feedback, and even more so when the feedback inhibition is delayed. When the excitation is delayed, the response correlations are instead increased compared to the no-feedback case for short time lags. These results thus show that cortical feedback may influence the temporal decorrelation of naturalistic stimuli, and that the degree of decorrelation depends on the spatiotemporal configuration of the feedback. In particular, our mixed feedback configuration seems to be particularly suited for reducing the temporal information redundancy in the signal.

## 4 Discussion

In the present work we have developed a firing-rate based simulation tool (`pyLGN`) to compute spatiotemporal responses of cells in the early visual system to visual stimuli. The simulation tool is based on the extended difference-of-gaussians (eDOG) model. This model provides closed-form expressions for (Fourier transformed) responses of both dLGN cells and cortical cells, also when cortical feedback projections to dLGN are explicitly included [40]. A main advantage of `pyLGN` is its computational and conceptual ease. The computation of visual responses corresponds to direct evaluation of two-dimensional or three-dimensional integrals in the case of static or dynamic (i.e., movie) stimuli, respectively, contrasting numerically extensive LGN network simulation based on spiking neurons [31, 35, 39, 82, 95] or models where each neuron is represented as individual firing-rate unit [36, 37]. This computational simplicity of `pyLGN` allows, for example, for fast and comprehensive exploration of a wide range of candidate scenarios for the organization of the cortical feedback.

### 4.1 Spatial effects of feedback

As a first example application we focused on the effect of cortical feedback on the spatial response properties of dLGN cells. A specific focus was on so-called area-response curves, i.e., responses to circular spots and patch-gratings as a function of stimulus size, which has received substantial experimental attention. In particular, studies have reported several effects of cortical feedback including sharpening of the receptive field by enhancing the center-surround antagonism of relay cells, increased receptive field center size with removal of feedback, and increased peak response to a an optimal diameter stimulus [16, 17, 22, 23, 96, 97]. Other studies have reported a more diverse influence from corticothalamic feedback, including both facilitatory and suppressive effects on dLGN cell responses, and changes in the receptive-field structure.

Our model demonstrated that cortical feedback can, depending on the feedback configuration, both enhance and suppress center-surround antagonism and both increase and decrease the receptive-field center size of relay cells. While the receptive-field center size decreases and the center-surround antagonism (as measured by the suppression index *α*_s_) increases with increased (indirect) cortical inhibitory feedback, the opposite is seen for excitatory feedback (Fig. 10).

These results support that a phase-reversed arrangement of the cortical feedback, where the ON-ON feedback is inhibitory while the OFF-ON feedback is excitatory, as suggested by data from [74], is more effective to enhance the center-surround antagonism of relay cells as observed in experiments [16, 17, 22, 23, 39]. However, with this arrangement a reduction in response to the optimal diameter stimulus in the size tuning curves was observed in our model (Fig. 10), in contrast to some experimental studies where an increase in response was reported [17, 96, 97]

Here we also considered the more complicated mixed phase-reversed feedback situation with a spatially broad ON-ON inhibitory feedback (combined with a corresponding OFF-ON excitatory feedback) and a spatially narrow ON-ON excitatory feedback (combined with a corresponding OFF-ON inhbitory feedback). Such a center-surround spatial organization of the feedback with excitatory bias to center and an inhibitory bias to the surround has been seen experimentally [3, 16]. In our model studies such mixed feedback was seen to give increased center-surround antagonism compared to the situation with ON-ON inhibitory feedback alone. Further, this configuration could also both reduce the size of the optimal stimulus diameter, as well as increase the magnitude of the response to the optimal stimulus diameter (Fig. 10). Correspondingly, with this configuration a sharper band-pass property of the spatial-frequency spectra was observed (Fig. 11).

### 4.2 Temporal effects of feedback

As for the spatial response properties, the effects of ON-ON inhibitory and ON-ON excitatory feedback (accompanied by the corresponding phase-reversed OFF-ON feedback) are seen to be quite distinct. While delayed inhibitory feedback makes the impulse response more biphasic, the opposite is the case for delayed excitatory feedback (Fig. 14). Likewise, while the temporal frequency tuning becomes sharper with delayed inhibitory feedback, it becomes blunter with delayed excitatory feedback (Fig. 16).

These features, i.e., increased biphasic index and sharper temporal frequency tuning, are maintained also for the case of mixed cortical feedback as long as the thalamocortical loop delay for the inhibitory feedback is much larger than for the excitatory feedback (Fig. 15, Fig. 17). Such a mixed-feedback configuration is also found to be particularly suited to remove temporal correlation in the stimulus and thus reduce the temporal redundancy in the neural signals that are sent from dLGN relay cells to cortex (Fig. 18).

### 4.3 Spatiotemporal organization of cortical feedback

Our results concerning the spatial and temporal feedback effects suggests that a situation with a mixed organization of cortical feedback consisting of a slow (long- delay) and spatially widespread ON-ON inhibitory feedback, combined with a fast (short-delay) and spatially narrow ON-ON excitatory feedback may have particular advantages. (Here the inhibitory and excitatory ON-ON feedback connections are accompanied by excitatory and inhibitory OFF-ON connections following a phase-reversed arrangement [39].)

This feedback organization seems well suited to dynamically modulate both the center-surround suppression and spatial resolution, for example, to adapt to changing light conditions where the most efficient neural representation of the stimulus is expected to vary depending on the signal-to-noise ratio [41]. In particular, for high light levels (i.e., high signal-to-noise) a band-pass like spatial spectrum (as obtained with our model for certain parameter choices) is expected to provide the most efficient coding, while for low light levels (low signal-to-noise) a low-pass spatial spectrum (as obtained with our model for some other parameter choices) seems better (see Sec. 3.6.1 in [98])

Further, a longer thalamocortical loop time of ON-ON inhibitory feedback compared to ON-ON excitatory feedback assures that temporal correlations in the natural visual stimuli are reduced in the relay-cell responses (Fig. 18). This temporal feedback arrangement gives a large biphasic index (Fig. 15) which previously has been shown to provide temporal decorrelation of natural stimuli [42], a feature that has also been seen in experiments [43]. Further, while the slow inhibitory feedback is key for providing this decorrelation, the rapid excitatory feedback may have a role in linking stimulus features by synchronising firing of neighbouring relay cells to provide a strong input to cortical target cells [10, 19]. Interestingly, a recent study found a large variation in axonal conduction times for corticothalamic axons, from a few milliseconds to many tens of milliseconds [99]. This suggests that differences in feedback delays indeed may have a functional role.

It should be noted that the eDOG-model [40] on which `pyLGN` is based, assumes that the cortical feedback has a phase-reversed arrangement where each ON-ON feedback connection is accompanied by a phase-reversed OFF-ON feedback connection (Eq. 11), i.e., a push-pull arrangement as experimentally observed in [74]. An alternative is a phase-matched arrangement where relay cells receive feedback only from cortical cells with the same symmetry, including both the direct excitatory feedback and the indirect inhibitory feedback. However, such an arrangement is not only at odds with the observations in [74], but also fails to explain the experimentally observed cortical-feedback induced increase in center-surround antagonism [39].

### 4.4 Outlook

Compared to the primary visual cortex (V1), i.e., the next station in the early visual pathway, the dLGN has received relatively little attention from computational neuroscientists [100]. From a modelling strategy point of view, this is somewhat unnatural as progress towards a mechanistic understanding of the function of the dLGN circuit seems more attainable given that (i) the dLGN circuit involves much fewer neuron types and is more comprehensively mapped out [2, 30], and that (ii) the dLGN has much fewer neurons making simulations computationally less intensive (18000 neurons in dLGN vs. 360000 neurons in V1 in mouse [101, 102]). Further, the strong recurrent interactions characteristic for cortical networks (which make them difficult to understand and analyse) appear absent between the principal cells (relay cells) in the dLGN, even if circuit network motifs such as feedforward and feedback interactions obviously are present. Thus a focused and comprehensive effort on mechanistic modeling of the dLGN circuit would not only be of interest in itself, it would also likely be a very useful stepping stone for later attempts to model the visual cortex.

While network simulations based on biophysically-detailed neuron models (e.g., [39, 82]) for entire dLGN nuclei are becoming computationally feasible with modern computers, there will still be a need for conceptually and mathematically simpler network simulation tools such as the present `pyLGN` tool based on the eDOG model. Such models are important to gain intuition about how the different circuit components may affect the overall circuit behaviour, and will also be important for guiding the choice of the numerous, typically unknown, parameters in more comprehensive dLGN network simulations. Thus we envision that a future mathematics-based understanding of the dLGN circuit will be of a ‘multiscale’ nature and be based on a set of interconnected models at different levels of biophysical detail. The mouse seems particularly suitable as model animal since construction and testing of the multiscale models can be greatly facilitated by the ever more sophisticated techniques for controlling gene expression in mice as well as the possibility for optogenetic activation [101, 103]).

We further envision that at a close and targeted collaboration between modellers and experimentalist, in particular direct assessment of predictions from mechanistic models in targeted experiments, holds great promise for unravelling mechanisms of visual information processing at the different levels of advancements in visual information processing from the earliest sensory systems to more complex computations of the higher cortical areas.

## Acknowledgments

We wish to thank Svenn-Arne Dragly, Simen Tenn e, and Helene Midtfjord for their helpful inputs and discussions regarding the design of the `pyLGN` simulator.

